# Insertion sequences associated with antibiotic resistance genes in *Enterococcus* isolates from an inpatient with prolonged bacteremia

**DOI:** 10.1101/2021.07.22.453447

**Authors:** Zulema Udaondo, Kaleb Z. Abram, Atul Kothari, Se-Ran Jun

## Abstract

Insertion sequences (ISs) and other transposable elements are associated with the mobilization of antibiotic resistance determinants and the modulation of pathogenic characteristics. In this work, we aimed to investigate the association between ISs and antibiotic resistance genes, and their role in dissemination and modification of antibiotic resistance phenotype. To that end, we leveraged fully resolved *Enterococcus faecium* and *Enterococcus faecalis* genomes of isolates collected over four days from an inpatient with prolonged bacteremia. Isolates from both species harbored similar IS family content but showed significant species-dependent differences in copy number and arrangements of ISs throughout their replicons. Here, we describe two inter-specific IS-mediated recombination events, and IS-medicated excision events in plasmids of *E. faecium* isolates. We also characterize a novel arrangement of the IS in a Tn1546-like transposon in *E. faecalis* isolates likely implicated in a vancomycin genotype-phenotype discrepancy. Furthermore, an extended analysis revealed a novel association between daptomycin resistance mutations in *liaSR* genes and a putative composite transposon in *E. faecium* offering a new paradigm for the study of daptomycin-resistance and novel insights into the route of daptomycin resistance dissemination. In conclusion, our study highlights the role ISs and other transposable elements play in rapid adaptation and response to clinically relevant stresses such as aggressive antibiotic treatment in enterococci.

## Introduction

*Enterococcus* spp. are ubiquitous bacteria that live as commensals in the gastrointestinal tract of humans and other mammals (1, 2). Their genetic plasticity enables them to acquire genetic determinants (3) to thrive in modern hospital environments where they often cause severe infectious diseases, especially in immunocompromised patients (2, 4). Multidrug-resistant enterococci not only play an important role as nosocomial pathogens, but they are a reservoir of genes encoding antibiotic resistance that can be transferred to other commensal microorganisms, increasing the risk of colonization and infection (4, 5).

In recent years, vancomycin-resistant *Enterococcus* (VRE) infection cases have increased rapidly around the world. As a consequence, a number of studies have examined the epidemiology (6, 7), population dynamics (8), and evolutionary landscape (9) of *Enterococcus*. However, there is still a lack of knowledge about the level of influence that plasmids and mobile genetic elements have on the persistence and spread of VRE strains (10). This is partly due to the presence of repetitive regions in genomes of *Enterococcus* that hinder full genome reconstruction (11, 12). Some of these repetitive sequences, such as insertion sequences (ISs), have been considered pathogenicity enhancers (5,13,14). Therefore, analyzing these elements is of vital importance to determine key characteristics of the VRE strains, such as level of antimicrobial resistance and virulence potential or to identify pathogenic variants.

ISs are small autonomous transposable elements, generally flanked by short terminal inverted repeats, that usually carry at least one transposase (Tn) gene whose product mediates the transposition process (Figure 1A) (15, 16). ISs are not only able to move themselves to different locations within genomes through various mechanisms (17) but are also able to travel horizontally between genomes as part of other mobile genetic elements such as bacteriophages and plasmids (15, 18). Transposases are mainly classified according to their transposition chemistry related to their catalytic domain topology and also based on their mode of transposition (replicative or non-replicative) (16–18). Furthermore, DNA segments carrying several genes can move together as a single unit when flanked by the same (or related) properly oriented ISs (15, 16). These larger transposable units are called compound or composite transposons (Figure 1A). Well-known examples of composite transposons that carry antibiotic resistance genes (ARGs) are Tn5 (kanamycin resistance) (19), Tn9 (chloramphenicol resistance) (20), and Tn10 (tetracycline resistance) (21). The important roles of ISs in transmission, inactivation, and modification of the resistant genotype and virulence of prokaryotic strains have been reviewed in previous studies (15, 18). However, most of these studies were performed using contigs or scaffolds from genome assemblies that might be missing a number of repetitive ISs.

**Figure 1.**
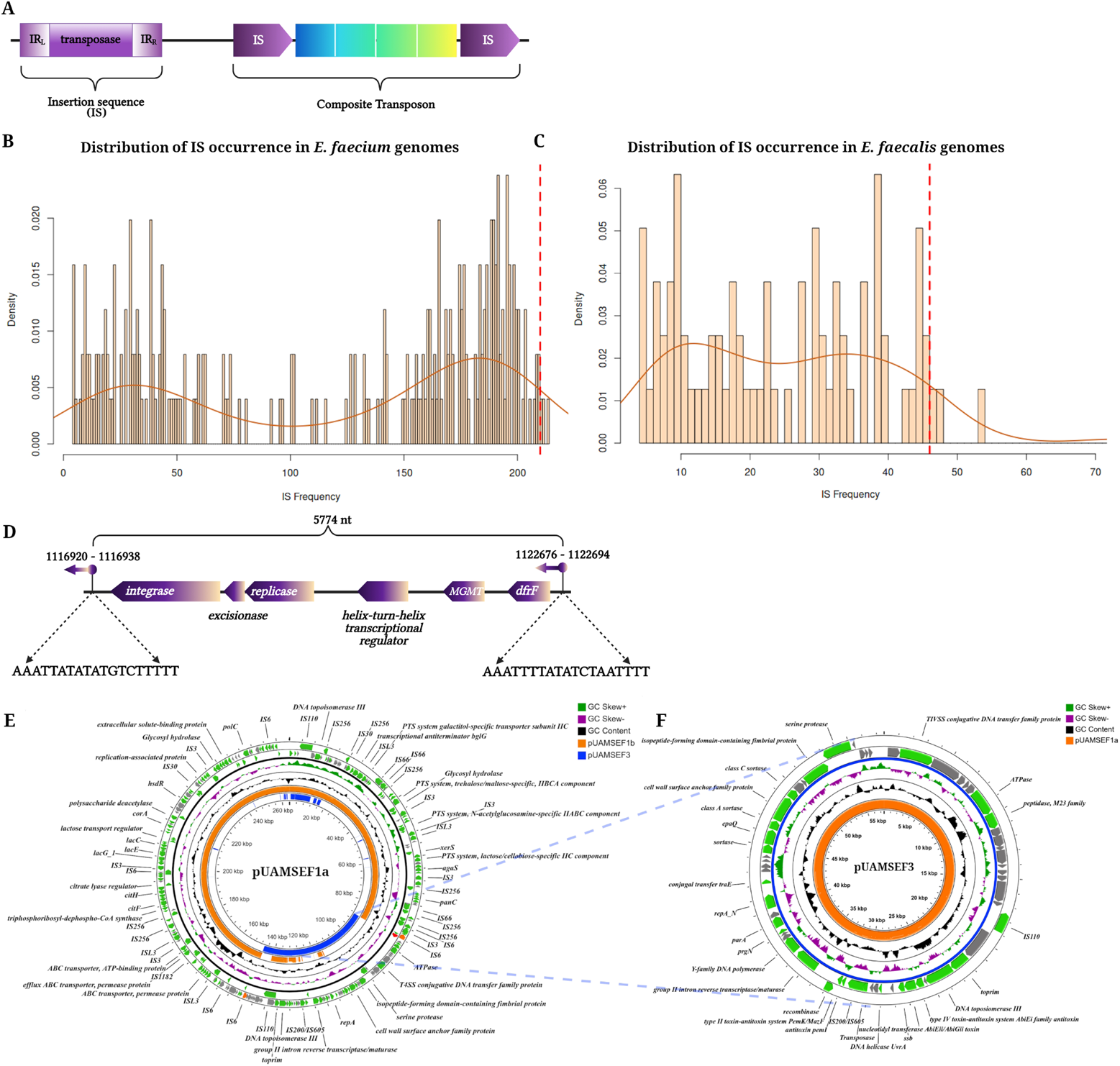
**A.** Schematic representations of an insertion sequence, in which the transposase consumes nearly the entire length of the element, and a composite transposon. The IS elements in the composite transposon are represented according to the orientation of their encoded transposase. Both IS elements can also be oriented inversely. **B and C.** Distribution of IS occurrence in (B) 252 *E. faecium* complete genome sequences and (C) 79 *E. faecalis* using ISEScan v1.7.2.1. Y-axis indicates the density of each IS frequency, where density is calculated as the number of occurrences for a given IS divided by the total number of genomes (252 for *E. faecium,* 79 for *E. faecalis*). X-axis indicates count of ISs present in genomes. The mean of the number of IS families annotated in the 3 *E. faecalis* isolates in this study is marked with a red dashed line. **D.** Schematic representation of novel transposable element found in *E. faecalis* isolates from this study. This figure was performed usingBioRender. **E.** Plasmids pUAMSEF1a from the first collected isolate of *E. faecium*, UAMS_EF55. **F.** Plasmid pUAMSEF3, a product of IS-mediated excision from pUAMSEF1a, present in UAMS_EF57 and UAMS_EF58 isolates.

In this study, we provide insights into the landscape of transposable elements that could be involved in disseminating multidrug resistance to bacteria in co-infection with VRE. For this purpose, we analyzed 6 bacterial isolates collected from the blood of an inpatient with bloodstream co-infection and diagnosed with Acute Lymphocytic Leukemia. The fully resolved high-quality genome assemblies of 6 isolates obtained from the patient (3 *Enterococcus faecium* and 3 *Enterococcus faecalis* isolates) were generated using Oxford Nanopore MinION and Illumina reads through a hybrid *de novo* strategy. Assemblies were screened for ARGs and mutations related to antimicrobial resistance, especially focusing on vancomycin and daptomycin resistance. Chromosomal and plasmid positions of repetitive ISs that usually flank bacterial ARGs and virulence genes were annotated. Our results shed light on the essential role ISs and other transposable elements play in large-scale genome variation due to IS-mediated genetic rearrangements and their association with ARGs in bacterial cells that persist in stressful environments under antibiotic exposure.

## Materials and Methods

### Sample collection and vancomycin susceptibility testing

The 6 VRE isolates were identified from positive blood cultures from one patient with prolonged bacteremia at the University of Arkansas for Medical Sciences (UAMS). Samples collected from the patient at different time points were grown on blood agar cultures and processed on the BacT/ALERT 3D system (bioMérieux, Durham, NC, USA). The Accelerate Pheno system (Accelerate Diagnostics, Tucson, AZ, USA) was used for identification and susceptibility testing of initial positive blood cultures. For repeat positives, the identification and susceptibility testing were performed with Vitek 2 MS and Vitek 2 systems. Vancomycin resistance was confirmed using E-tests (bioMérieux), and antimicrobial susceptibility test results were interpreted using the M100 CLSI standards (22).

### Whole-genome sequencing

Genomic DNA was extracted and sequenced as described in (23, 24). Briefly, genomic DNA was extracted from VRE colonies using the Quick-DNA Fungal/Bacterial kit (Zymo Research, Irvine, CA, USA). A NanoDrop Spectrophotometer (Thermo Fisher, Waltham, MA, USA) was used to determine the purity of the extracted DNA by measuring the A260/280 and A260/230 ratios. Integrity and quantity of DNA were determined using an Agilent 2200 TapeStation (Agilent, Santa Clara, CA, USA) and Qubit 3.0 Assay (Thermo Fisher), respectively. Purified DNA was aliquoted into 2 tubes for MinION (Oxford Nanopore Technologies, Oxford, UK) and Illumina (San Diego, CA, USA) sequencing. Oxford Nanopore Technologies (ONT) sequencing libraries were prepared using a PCR-free method of multiplexing samples with the Rapid Barcoding kit (SQK-RBK004, ONT), and sequencing of the barcoded DNA was performed on a single R9.4/FLO-MIN106 ONT flow cell on the MinION (version Mk1B, ONT) for 48 hours. Illumina libraries for the 6 isolates were also sequenced using the NextSeq 550 platform at the UAMS Myeloma Center. Adapters were trimmed using fastp, version 0.19.5 (25), with default settings. Trimmomatic, version 0.38 (26), was used to remove poor-quality reads. The quality of pre- and post-processed reads was assessed with the FastQC tool, version 0.11.8 (27).

### Base calling and assembling

Base calling of Nanopore reads was performed using Guppy v4.4.2 (28), with -- min_qscore 8. Libraries were demultiplexed using guppy_barcoder from Guppy v4.4.2. Porechop v0.2.3 (https://github.com/rrwick/Porechop) was used to remove adapters, and NanoFilt from NanoPack (29) was used for further quality filtering. Read quality was examined with NanoStat v1.2.0 from NanoPack (29). Two different strategies were established to obtain fully resolved assemblies depending on the species. For *E. faecium* isolates, we used Unicycler v0.4.8 (30) hybrid assembler, with Canu v2.1.1 (31)-generated scaffolds and Illumina data as inputs. Fully resolved assemblies for *E. faecalis* isolates were obtained using Flye v2.8.2 assembler (32) with Oxford Nanopore reads, with 2 rounds of polishing using the Oxford Nanopore long-reads with Racon v1.4.20 (33) and Medaka v1.2.1 (https://nanoporetech.github.io/medaka/index.html). For both *E. faecium* and *E. faecalis*, final polishing with Illumina data was performed through 2 rounds of Pilon v1.23 (34).

### Functional antimicrobial resistance and mobilome annotation

Functional annotation of the 6 isolates was performed using Prokka v1.14.6 (35) with the --rnammer option. Functional annotation of plasmid replicons and other chromosomal regions of interest were manually reannotated using PsiBlast and Blastp from NCBI and Enterococcus nr/nt database. Antimicrobial resistance genes were annotated with Resistance Gene Identifier (RGI) software against the Comprehensive Antibiotic Resistance Database (CARD) (36). ISEScan v1.7.2.1 (37) (Xie and Tang, 2017) and ISfinder database (https://isfinder.biotoul.fr/) (38) were used to identify transposable elements and their inverted-repeat sequences. Plasmids were classified using PlasmidFinder2.0 (39). CGview server (version beta) (40) was used to plot functional and structural characteristics of the plasmids in this study.

### Multilocus sequence typing (MLST)

MLST analysis was performed using mlst tool (Seemann T, `mlst` Github https://github.com/tseemann/mlst) with public PubMLST typing schemes (41).

### Comparative genome analysis

All replicons were divided into fragments of 5000 bp with an overlap of 3000 bp (2000 step size) and compared via Blastn v2.9.0 (42). Bidirectional best hits between replicons from different isolates were identified using best bit score, e-value, and percentage of sequence similarity. Mash software v2.3 (43) with k=21 was used to determine genetic distances between isolates where sketch option with −s 10000 was used to increase the sensitivity. Snippy (https://github.com/tseemann/snippy) was used to analyze the coreSNP of the 6 isolates by species affiliation.

Figures were prepared using Clinker v0.0.20 (44), pyGenomeTracks (45) (Lopez-Delisle et al., 2021), and BioRender (https://biorender.com/).

## Results

### Fully resolved replicons in *E. faecalis* and *E. faecium* isolates obtained from hybrid *de novo* assembly

The 6 isolates from one patient with prolonged bacteremia were collected from positive blood cultures over a period of 4 days and identified as *E. faecalis* (isolates UAMS_EL53, UAMS_EL54, and UAMS_EL56) or *E. faecium* (UAMS_EF55, UAMS_EF57, and UAMS_EF58). Vancomycin susceptibility testing was negative for all *E. faecalis* isolates (minimum inhibitory concentration, MIC ≤ 2 μg/ml) indicating a susceptible phenotype. However, all *E. faecium* isolates were phenotypically resistant to vancomycin, with MIC > 16 μg/ml for the first 2 isolates and MIC ≥ 64 μg/mL for the last collected isolate (UAMS_EF58). Two different *de novo* hybrid assembly strategies, depending on species (described in Materials and Methods) were used to generate fully resolved assemblies for all 6 isolates. Thus, complete circular replicons were obtained for the 6 chromosomes and for almost all of the plasmids for the 6 isolates in this study (Table 1).

**Table 1.** Summary of *de novo* assembly results for *E. faecalis* and *E. faecium* isolates.

| Isolate | # | Species | Chromosome | Plasmid 1 | Plasmid 2 | Plasmid 3 | Plasmid 4 | Plasmid 5 |
| --- | --- | --- | --- | --- | --- | --- | --- | --- |
| UAMS_EL53 | 1 | *E. faecalis* | 3 030 005c | 119 720c | 55 301c | 53 420c | 33 652c x3 |  |
| UAMS_EL54 | 2 | *E. faecalis* | 3 030 422c | 119 714c | 55 304c | 53 047c | 33 656c x3 |  |
| UAMS_EL56 | 4 | *E. faecalis* | 3 031 832c | 119 713c | 55 288c | 53 193c | 33 659c x3 |  |
| UAMS_EF55 | 3 | *E. faecium* | 2 841 245c | 264 192c | 79 782 |  | 41 819c x3 | 4 374c |
| UAMS_EF57 | 5 | *E. faecium* | 2 841 244c | 205 436c | 79 769 | 57 509c | 41 819c x3 | 4 374c |
| UAMS_EF58 | 6 | *E. faecium* | 2 841 249c | 205 407c | 79 679 | 57 509c | 34 693c x3 | 4 374c |

Lengths (bp) of replicons found in each isolate are shown. #: Order by sample collection time. c: Indicates circularized replicon according to Assembler. x3: Indicates inferred multiplicity based on normalized read depth per contig. Plasmid 1, UAMS\_EF55 = pUAMSEF1a; Plasmid 1, UAMS\_EF57 = pUAMSEF1b; and Plasmid 1, UAMS\_EF58 = pUAMSEF1b, in the main text.

As expected, chromosome sizes for *E. faecalis* isolates (classified as ST6 by multilocus sequence typing, MLST) were slightly larger (∼3 Mb) than those in *E. faecium* (classified as ST736 by MLST) isolates (∼2.8 Mb). Four to 5 complete plasmids were found in all isolates (Table 1). All 6 isolates harbored plasmids with a complete set of genes conferring vancomycin resistance (*vanR*, *vanS*, *vanH*, *vanA*, *vanX*, and *vanZ*; VanA phenotype), revealing a genotypic– phenotypic discrepancy in *E. faecalis* isolates. It is worth noting that plasmids carrying vancomycin resistance genes presented greater differences in read depth compared to the other replicons in this study. The multiplicity of *vanA* replicons was inferred to be triplicated in all assemblies of the study based on the normalized read depth (Table 1). Plasmid genomes comparison analysis showed that *vanA* plasmid from *E. fecalis* are most similar to *vanA* plasmid from *E. faecium* (Suppl. Figure 1A). Mash analysis of whole genome assemblies performed on the 6 isolates of the study showed that the last 2 collected isolates for each species had the largest intra-specific variability (Suppl. Figure 1B). Mash distances between isolates from different species were about ∼0.18.

### Arrangement, copy number, and association of insertion sequences with antibiotic resistance genes differ in *E. faecalis* and *E. faecium* isolates

Detailed analysis of the ISs annotated by ISEScan (37) and ISfinder (38) in our 6 isolates identified 11 different IS families in *E. faecalis* isolates, including one putative novel IS family, and 12 IS families in *E. faecium* isolates (Table 2). Copy number of the IS families identified in both species varied considerably, with these elements being 5 times more abundant in *E. faecium* isolates (∼210 IS/isolate) than in *E. faecalis* isolates (∼46 IS/isolate). In *E. faecalis* isolates, the IS*256* family (n=11 in the first two collected isolates and n=12 in the last collected isolate) was the most abundant IS, followed by ISs from IS*6 (*n*=*9 in each isolate) and IS*3* families (n=9 in each isolate). In *E. faecium* isolates, the IS*L3* family (n=61) was the most abundant, with elements distributed mainly throughout the chromosome, followed by elements from families IS*256* (n=35) and IS*30* (n=27). Sequences from IS*66* and IS*1380* families were found in all *E. faecium* isolates but not in *E. faecalis*. The IS elements identified in isolates from both species were distributed evenly along the chromosome (Table 2), but they were also found distributed among 3 of the 4 identified plasmids in *E. faecalis* isolates and in 4 of the 5 plasmids in *E. faecium*. Copy number of the annotated ISs was preserved in all 3 *E. faecalis* isolates with the exception of one additional element from the IS*256* family identified in the chromosome of UAMS_EL56. This IS*256* element was located in an area surrounded by other ISs from IS*256* and IS*6* families, in which the only characterized gene was choloylglycine hydrolase family protein (Suppl. Table 1).

**Table 2.**
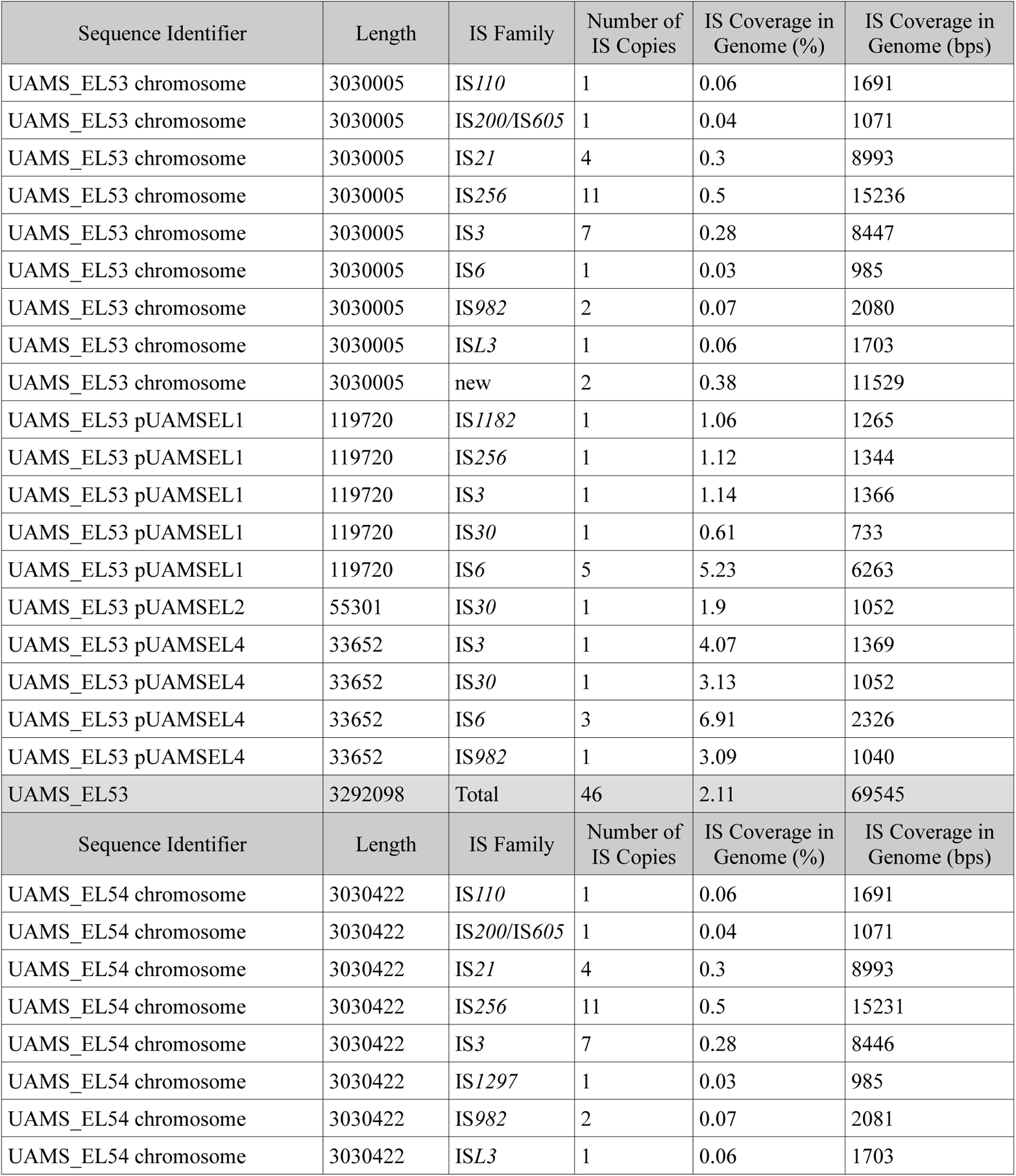
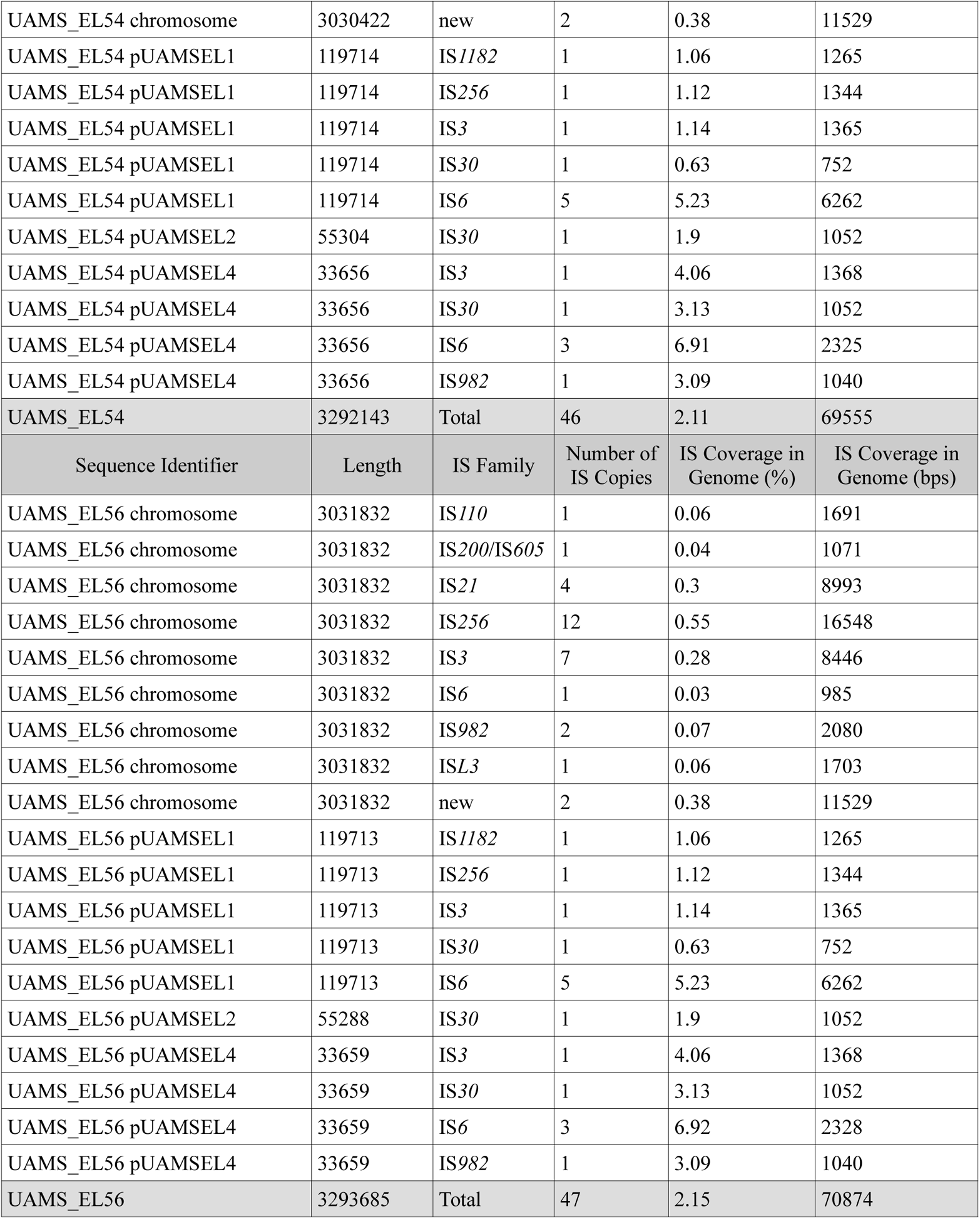
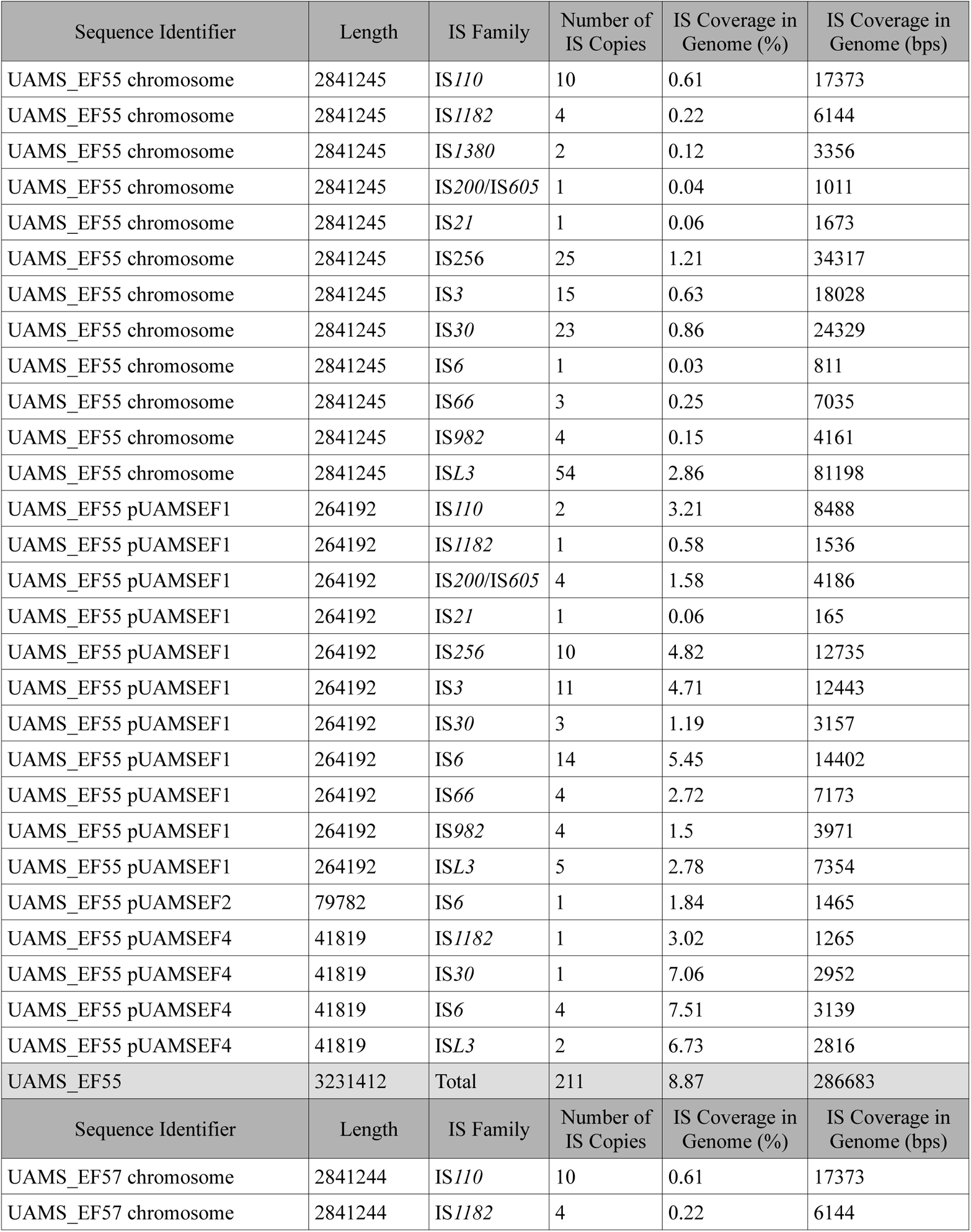
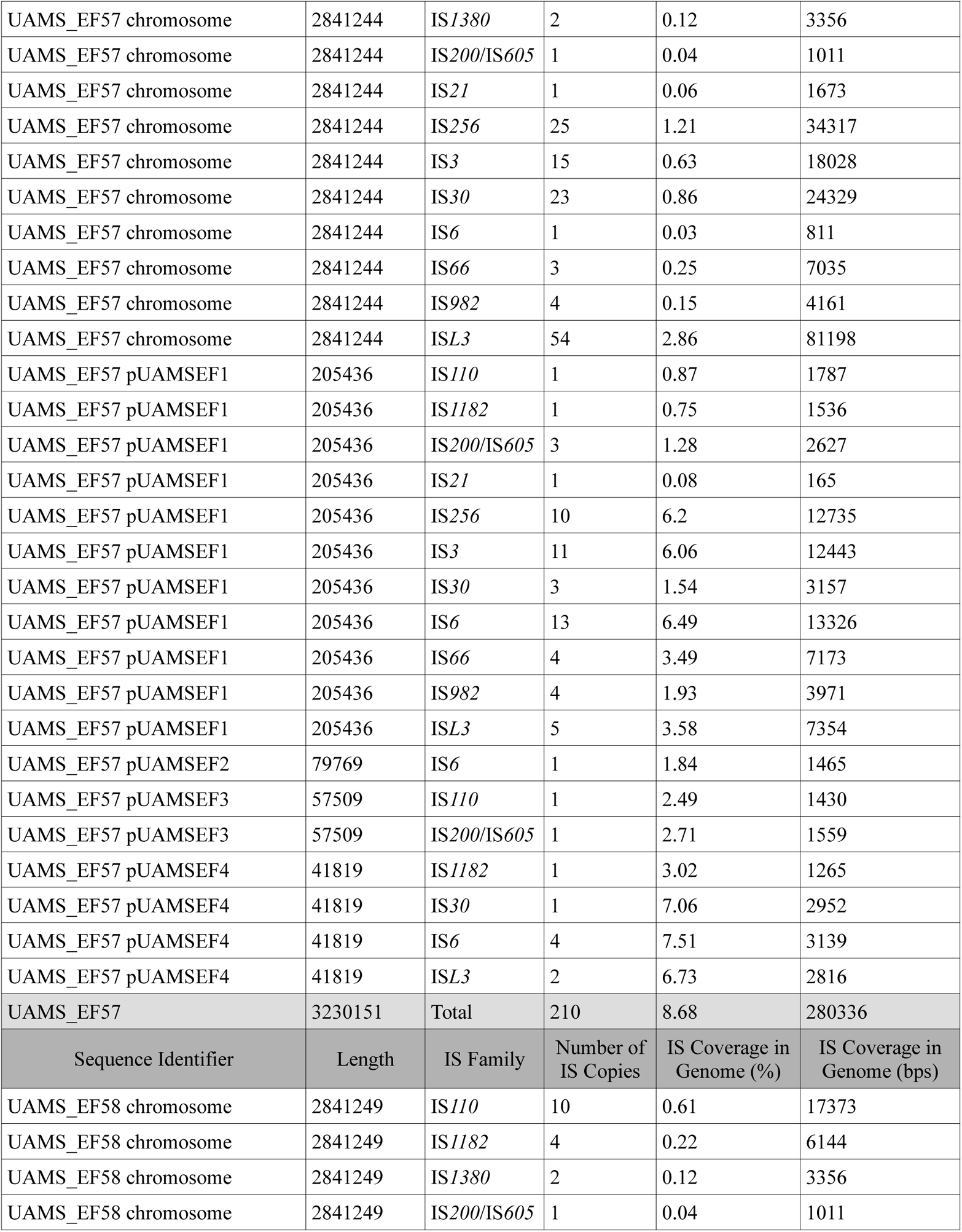
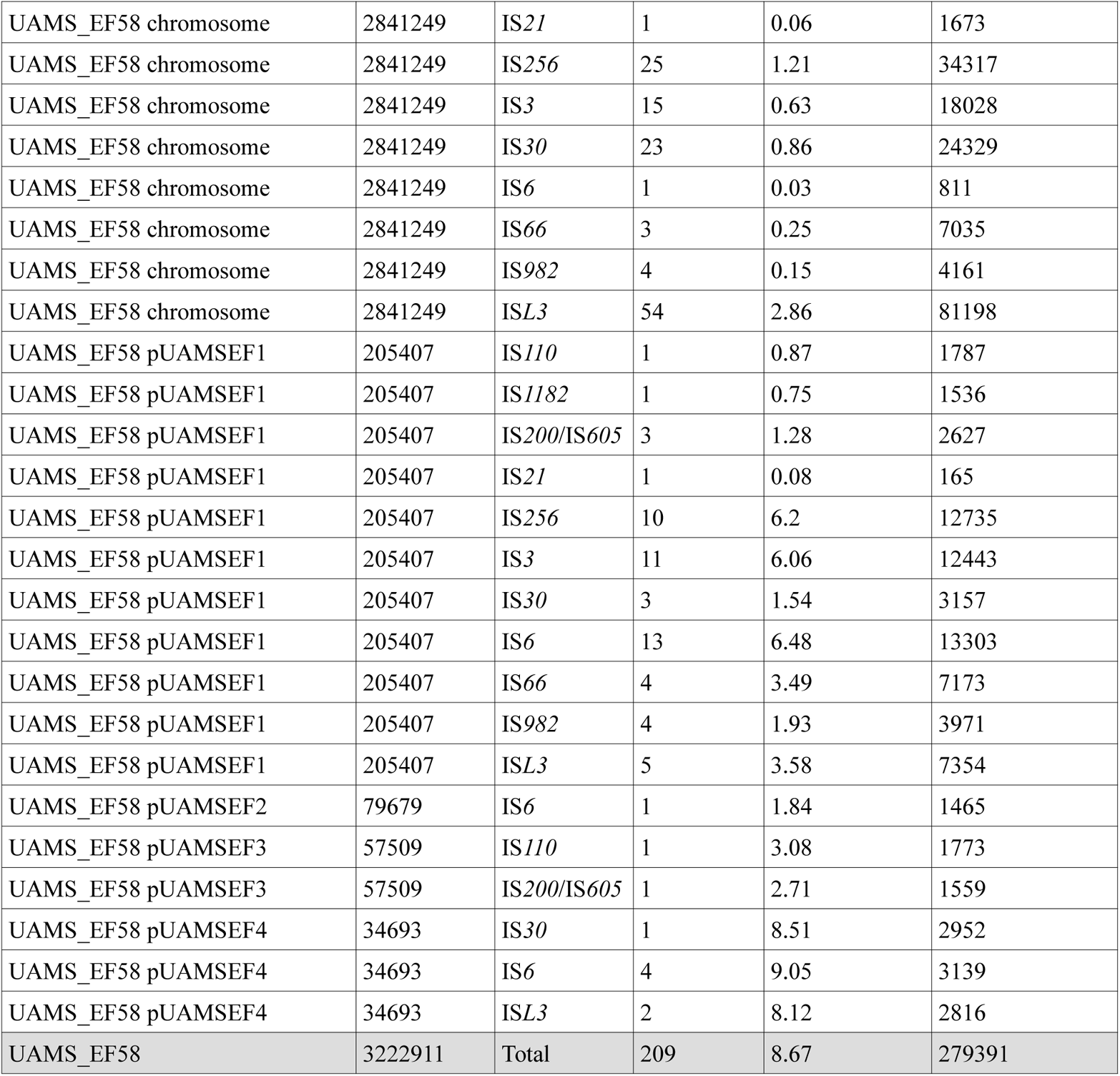
Summary of annotated insertion sequences in each of the *E. faecalis* and *E. faecium* isolates using ISEScan.

| Sequence Identifier | Length | IS Family | Number of IS Copies | IS Coverage in Genome (%) | IS Coverage in Genome (bps) |
| --- | --- | --- | --- | --- | --- |
| UAMS_EL53 chromosome | 3030005 | IS110 | 1 | 0.06 | 1691 |
| UAMS_EL53 chromosome | 3030005 | IS200/IS605 | 1 | 0.04 | 1071 |
| UAMS_EL53 chromosome | 3030005 | IS21 | 4 | 0.3 | 8993 |
| UAMS_EL53 chromosome | 3030005 | IS256 | 11 | 0.5 | 15236 |
| UAMS_EL53 chromosome | 3030005 | IS3 | 7 | 0.28 | 8447 |
| UAMS_EL53 chromosome | 3030005 | IS6 | 1 | 0.03 | 985 |
| UAMS_EL53 chromosome | 3030005 | IS982 | 2 | 0.07 | 2080 |
| UAMS_EL53 chromosome | 3030005 | ISL3 | 1 | 0.06 | 1703 |
| UAMS_EL53 chromosome | 3030005 | new | 2 | 0.38 | 11529 |
| UAMS_EL53 pUAMSEL1 | 119720 | IS1182 | 1 | 1.06 | 1265 |
| UAMS_EL53 pUAMSEL1 | 119720 | IS256 | 1 | 1.12 | 1344 |
| UAMS_EL53 pUAMSEL1 | 119720 | IS3 | 1 | 1.14 | 1366 |
| UAMS_EL53 pUAMSEL1 | 119720 | IS30 | 1 | 0.61 | 733 |
| UAMS_EL53 pUAMSEL1 | 119720 | IS6 | 5 | 5.23 | 6263 |
| UAMS_EL53 pUAMSEL2 | 55301 | IS30 | 1 | 1.9 | 1052 |
| UAMS_EL53 pUAMSEL4 | 33652 | IS3 | 1 | 4.07 | 1369 |
| UAMS_EL53 pUAMSEL4 | 33652 | IS30 | 1 | 3.13 | 1052 |
| UAMS_EL53 pUAMSEL4 | 33652 | IS6 | 3 | 6.91 | 2326 |
| UAMS_EL53 pUAMSEL4 | 33652 | IS982 | 1 | 3.09 | 1040 |
| UAMS_EL53 | 3292098 | Total | 46 | 2.11 | 69545 |
| Sequence Identifier | Length | IS Family | Number of IS Copies | IS Coverage in Genome (%) | IS Coverage in Genome (bps) |
| UAMS_EL54 chromosome | 3030422 | IS110 | 1 | 0.06 | 1691 |
| UAMS_EL54 chromosome | 3030422 | IS200/IS605 | 1 | 0.04 | 1071 |
| UAMS_EL54 chromosome | 3030422 | IS21 | 4 | 0.3 | 8993 |
| UAMS_EL54 chromosome | 3030422 | IS256 | 11 | 0.5 | 15231 |
| UAMS_EL54 chromosome | 3030422 | IS3 | 7 | 0.28 | 8446 |
| UAMS_EL54 chromosome | 3030422 | IS1297 | 1 | 0.03 | 985 |
| UAMS_EL54 chromosome | 3030422 | IS982 | 2 | 0.07 | 2081 |
| UAMS_EL54 chromosome | 3030422 | ISL3 | 1 | 0.06 | 1703 |

| UAMS_EL54 chromosome | 3030422 | new | 2 | 0.38 | 11529 |
| --- | --- | --- | --- | --- | --- |
| UAMS_EL54 pUAMSEL1 | 119714 | IS1182 | 1 | 1.06 | 1265 |
| UAMS_EL54 pUAMSEL1 | 119714 | IS256 | 1 | 1.12 | 1344 |
| UAMS_EL54 pUAMSEL1 | 119714 | IS3 | 1 | 1.14 | 1365 |
| UAMS_EL54 pUAMSEL1 | 119714 | IS30 | 1 | 0.63 | 752 |
| UAMS_EL54 pUAMSEL1 | 119714 | IS6 | 5 | 5.23 | 6262 |
| UAMS_EL54 pUAMSEL2 | 55304 | IS30 | 1 | 1.9 | 1052 |
| UAMS_EL54 pUAMSEL4 | 33656 | IS3 | 1 | 4.06 | 1368 |
| UAMS_EL54 pUAMSEL4 | 33656 | IS30 | 1 | 3.13 | 1052 |
| UAMS_EL54 pUAMSEL4 | 33656 | IS6 | 3 | 6.91 | 2325 |
| UAMS_EL54 pUAMSEL4 | 33656 | IS982 | 1 | 3.09 | 1040 |
| UAMS_EL54 | 3292143 | Total | 46 | 2.11 | 69555 |
| Sequence Identifier | Length | IS Family | Number of IS Copies | IS Coverage in Genome (%) | IS Coverage in Genome (bps) |
| UAMS_EL56 chromosome | 3031832 | IS110 | 1 | 0.06 | 1691 |
| UAMS_EL56 chromosome | 3031832 | IS200/IS605 | 1 | 0.04 | 1071 |
| UAMS_EL56 chromosome | 3031832 | IS21 | 4 | 0.3 | 8993 |
| UAMS_EL56 chromosome | 3031832 | IS256 | 12 | 0.55 | 16548 |
| UAMS_EL56 chromosome | 3031832 | IS3 | 7 | 0.28 | 8446 |
| UAMS_EL56 chromosome | 3031832 | IS6 | 1 | 0.03 | 985 |
| UAMS_EL56 chromosome | 3031832 | IS982 | 2 | 0.07 | 2080 |
| UAMS_EL56 chromosome | 3031832 | ISL3 | 1 | 0.06 | 1703 |
| UAMS_EL56 chromosome | 3031832 | new | 2 | 0.38 | 11529 |
| UAMS_EL56 pUAMSEL1 | 119713 | IS1182 | 1 | 1.06 | 1265 |
| UAMS_EL56 pUAMSEL1 | 119713 | IS256 | 1 | 1.12 | 1344 |
| UAMS_EL56 pUAMSEL1 | 119713 | IS3 | 1 | 1.14 | 1365 |
| UAMS_EL56 pUAMSEL1 | 119713 | IS30 | 1 | 0.63 | 752 |
| UAMS_EL56 pUAMSEL1 | 119713 | IS6 | 5 | 5.23 | 6262 |
| UAMS_EL56 pUAMSEL2 | 55288 | IS30 | 1 | 1.9 | 1052 |
| UAMS_EL56 pUAMSEL4 | 33659 | IS3 | 1 | 4.06 | 1368 |
| UAMS_EL56 pUAMSEL4 | 33659 | IS30 | 1 | 3.13 | 1052 |
| UAMS_EL56 pUAMSEL4 | 33659 | IS6 | 3 | 6.92 | 2328 |
| UAMS_EL56 pUAMSEL4 | 33659 | IS982 | 1 | 3.09 | 1040 |
| UAMS_EL56 | 3293685 | Total | 47 | 2.15 | 70874 |
| UAMS_EF55 chromosome | 2841245 | *IS110* | 10 | 0.61 | 17373 |
| UAMS_EF55 chromosome | 2841245 | *IS1182* | 4 | 0.22 | 6144 |
| UAMS_EF55 chromosome | 2841245 | *IS1380* | 2 | 0.12 | 3356 |
| UAMS_EF55 chromosome | 2841245 | *IS200/IS605* | 1 | 0.04 | 1011 |
| UAMS_EF55 chromosome | 2841245 | *IS21* | 1 | 0.06 | 1673 |
| UAMS_EF55 chromosome | 2841245 | *IS256* | 25 | 1.21 | 34317 |
| UAMS_EF55 chromosome | 2841245 | *IS3* | 15 | 0.63 | 18028 |
| UAMS_EF55 chromosome | 2841245 | *IS30* | 23 | 0.86 | 24329 |
| UAMS_EF55 chromosome | 2841245 | *IS6* | 1 | 0.03 | 811 |
| UAMS_EF55 chromosome | 2841245 | *IS66* | 3 | 0.25 | 7035 |
| UAMS_EF55 chromosome | 2841245 | *IS982* | 4 | 0.15 | 4161 |
| UAMS_EF55 chromosome | 2841245 | *ISL3* | 54 | 2.86 | 81198 |
| UAMS_EF55 pUAMSEF1 | 264192 | *IS110* | 2 | 3.21 | 8488 |
| UAMS_EF55 pUAMSEF1 | 264192 | *IS1182* | 1 | 0.58 | 1536 |
| UAMS_EF55 pUAMSEF1 | 264192 | *IS200/IS605* | 4 | 1.58 | 4186 |
| UAMS_EF55 pUAMSEF1 | 264192 | *IS21* | 1 | 0.06 | 165 |
| UAMS_EF55 pUAMSEF1 | 264192 | *IS256* | 10 | 4.82 | 12735 |
| UAMS_EF55 pUAMSEF1 | 264192 | *IS3* | 11 | 4.71 | 12443 |
| UAMS_EF55 pUAMSEF1 | 264192 | *IS30* | 3 | 1.19 | 3157 |
| UAMS_EF55 pUAMSEF1 | 264192 | *IS6* | 14 | 5.45 | 14402 |
| UAMS_EF55 pUAMSEF1 | 264192 | *IS66* | 4 | 2.72 | 7173 |
| UAMS_EF55 pUAMSEF1 | 264192 | *IS982* | 4 | 1.5 | 3971 |
| UAMS_EF55 pUAMSEF1 | 264192 | *ISL3* | 5 | 2.78 | 7354 |
| UAMS_EF55 pUAMSEF2 | 79782 | *IS6* | 1 | 1.84 | 1465 |
| UAMS_EF55 pUAMSEF4 | 41819 | *IS1182* | 1 | 3.02 | 1265 |
| UAMS_EF55 pUAMSEF4 | 41819 | *IS30* | 1 | 7.06 | 2952 |
| UAMS_EF55 pUAMSEF4 | 41819 | *IS6* | 4 | 7.51 | 3139 |
| UAMS_EF55 pUAMSEF4 | 41819 | *ISL3* | 2 | 6.73 | 2816 |
| UAMS_EF55 | 3231412 | Total | 211 | 8.87 | 286683 |
| Sequence Identifier | Length | IS Family | Number of IS Copies | IS Coverage in Genome (%) | IS Coverage in Genome (bps) |
| UAMS_EF57 chromosome | 2841244 | *IS110* | 10 | 0.61 | 17373 |
| UAMS_EF57 chromosome | 2841244 | *IS1182* | 4 | 0.22 | 6144 |

| UAMS_EF57 chromosome | 2841244 | IS1380 | 2 | 0.12 | 3356 |
| --- | --- | --- | --- | --- | --- |
| UAMS_EF57 chromosome | 2841244 | IS200/IS605 | 1 | 0.04 | 1011 |
| UAMS_EF57 chromosome | 2841244 | IS21 | 1 | 0.06 | 1673 |
| UAMS_EF57 chromosome | 2841244 | IS256 | 25 | 1.21 | 34317 |
| UAMS_EF57 chromosome | 2841244 | IS3 | 15 | 0.63 | 18028 |
| UAMS_EF57 chromosome | 2841244 | IS30 | 23 | 0.86 | 24329 |
| UAMS_EF57 chromosome | 2841244 | IS6 | 1 | 0.03 | 811 |
| UAMS_EF57 chromosome | 2841244 | IS66 | 3 | 0.25 | 7035 |
| UAMS_EF57 chromosome | 2841244 | IS982 | 4 | 0.15 | 4161 |
| UAMS_EF57 chromosome | 2841244 | ISL3 | 54 | 2.86 | 81198 |
| UAMS_EF57 pUAMSEF1 | 205436 | IS110 | 1 | 0.87 | 1787 |
| UAMS_EF57 pUAMSEF1 | 205436 | IS1182 | 1 | 0.75 | 1536 |
| UAMS_EF57 pUAMSEF1 | 205436 | IS200/IS605 | 3 | 1.28 | 2627 |
| UAMS_EF57 pUAMSEF1 | 205436 | IS21 | 1 | 0.08 | 165 |
| UAMS_EF57 pUAMSEF1 | 205436 | IS256 | 10 | 6.2 | 12735 |
| UAMS_EF57 pUAMSEF1 | 205436 | IS3 | 11 | 6.06 | 12443 |
| UAMS_EF57 pUAMSEF1 | 205436 | IS30 | 3 | 1.54 | 3157 |
| UAMS_EF57 pUAMSEF1 | 205436 | IS6 | 13 | 6.49 | 13326 |
| UAMS_EF57 pUAMSEF1 | 205436 | IS66 | 4 | 3.49 | 7173 |
| UAMS_EF57 pUAMSEF1 | 205436 | IS982 | 4 | 1.93 | 3971 |
| UAMS_EF57 pUAMSEF1 | 205436 | ISL3 | 5 | 3.58 | 7354 |
| UAMS_EF57 pUAMSEF2 | 79769 | IS6 | 1 | 1.84 | 1465 |
| UAMS_EF57 pUAMSEF3 | 57509 | IS110 | 1 | 2.49 | 1430 |
| UAMS_EF57 pUAMSEF3 | 57509 | IS200/IS605 | 1 | 2.71 | 1559 |
| UAMS_EF57 pUAMSEF4 | 41819 | IS1182 | 1 | 3.02 | 1265 |
| UAMS_EF57 pUAMSEF4 | 41819 | IS30 | 1 | 7.06 | 2952 |
| UAMS_EF57 pUAMSEF4 | 41819 | IS6 | 4 | 7.51 | 3139 |
| UAMS_EF57 pUAMSEF4 | 41819 | ISL3 | 2 | 6.73 | 2816 |
| UAMS_EF57 | 3230151 | Total | 210 | 8.68 | 280336 |
| Sequence Identifier | Length | IS Family | Number of IS Copies | IS Coverage in Genome (%) | IS Coverage in Genome (bps) |
| UAMS_EF58 chromosome | 2841249 | IS110 | 10 | 0.61 | 17373 |
| UAMS_EF58 chromosome | 2841249 | IS1182 | 4 | 0.22 | 6144 |
| UAMS_EF58 chromosome | 2841249 | IS1380 | 2 | 0.12 | 3356 |
| UAMS_EF58 chromosome | 2841249 | IS200/IS605 | 1 | 0.04 | 1011 |
| UAMS_EF58 chromosome | 2841249 | IS21 | 1 | 0.06 | 1673 |
| UAMS_EF58 chromosome | 2841249 | IS256 | 25 | 1.21 | 34317 |
| UAMS_EF58 chromosome | 2841249 | IS3 | 15 | 0.63 | 18028 |
| UAMS_EF58 chromosome | 2841249 | IS30 | 23 | 0.86 | 24329 |
| UAMS_EF58 chromosome | 2841249 | IS6 | 1 | 0.03 | 811 |
| UAMS_EF58 chromosome | 2841249 | IS66 | 3 | 0.25 | 7035 |
| UAMS_EF58 chromosome | 2841249 | IS982 | 4 | 0.15 | 4161 |
| UAMS_EF58 chromosome | 2841249 | ISL3 | 54 | 2.86 | 81198 |
| UAMS_EF58 pUAMSEF1 | 205407 | IS110 | 1 | 0.87 | 1787 |
| UAMS_EF58 pUAMSEF1 | 205407 | IS1182 | 1 | 0.75 | 1536 |
| UAMS_EF58 pUAMSEF1 | 205407 | IS200/IS605 | 3 | 1.28 | 2627 |
| UAMS_EF58 pUAMSEF1 | 205407 | IS21 | 1 | 0.08 | 165 |
| UAMS_EF58 pUAMSEF1 | 205407 | IS256 | 10 | 6.2 | 12735 |
| UAMS_EF58 pUAMSEF1 | 205407 | IS3 | 11 | 6.06 | 12443 |
| UAMS_EF58 pUAMSEF1 | 205407 | IS30 | 3 | 1.54 | 3157 |
| UAMS_EF58 pUAMSEF1 | 205407 | IS6 | 13 | 6.48 | 13303 |
| UAMS_EF58 pUAMSEF1 | 205407 | IS66 | 4 | 3.49 | 7173 |
| UAMS_EF58 pUAMSEF1 | 205407 | IS982 | 4 | 1.93 | 3971 |
| UAMS_EF58 pUAMSEF1 | 205407 | ISL3 | 5 | 3.58 | 7354 |
| UAMS_EF58 pUAMSEF2 | 79679 | IS6 | 1 | 1.84 | 1465 |
| UAMS_EF58 pUAMSEF3 | 57509 | IS110 | 1 | 3.08 | 1773 |
| UAMS_EF58 pUAMSEF3 | 57509 | IS200/IS605 | 1 | 2.71 | 1559 |
| UAMS_EF58 pUAMSEF4 | 34693 | IS30 | 1 | 8.51 | 2952 |
| UAMS_EF58 pUAMSEF4 | 34693 | IS6 | 4 | 9.05 | 3139 |
| UAMS_EF58 pUAMSEF4 | 34693 | ISL3 | 2 | 8.12 | 2816 |
| UAMS_EF58 | 3222911 | Total | 209 | 8.67 | 279391 |

Genetic variability can be useful to withstand stressful conditions (18, 46). Rates of IS transpositions can be increased by environmental factors, such as changes in the host environment and presence of subinhibitory concentrations of antibiotics that serve as a signal to promote genetic variability (47). Thus, the strikingly high number of ISs found in our *E. faecium* isolates could be a consequence of high rates of replicative transposition due to the stressful and competitive conditions of the bloodstream environment, which is not the natural habitat of enterococci. To place our isolates into a global context, we downloaded all complete *E. faecalis* and *E. faecium* genomes from GenBank (in Dec. 2020) and annotated them using the same pipeline used for the 6 isolates in this study. Figure 1B indicates the distribution of the number of IS families annotated in 252 complete *E. faecium* genome assemblies where the mean of the number of IS families annotated in our 3 *E. faecium* isolates is shown as an example of an extreme case of IS replicative transposition (red line, Figure 1B). The same scenario was observed in *E. faecalis*, where only three of 79 complete genome assemblies had more IS families than our clinical isolates (Figure 1C). Both species showed a bimodal distribution of annotated ISs, this distribution being accentuated in *E. faecium* strains, where most of the annotated genomes had a relatively small number of annotated ISs (n ≤ 50) or over 150 annotated ISs (n ≥ 190). However, we could not recover metadata related to the environmental source of most of the strains in this analysis, which prevented us from establishing correlation between the number of annotated ISs and the environmental source.

Considering a genetic neighborhood of 10 upstream and downstream annotated open reading frames (ORFs), 14 out of 21 ARGs were found in the proximity of at least one IS in our 3 *E. faecalis* isolates (Suppl. Table 1). Further analysis of the possible association between ISs and ARGs identified a previously unknown IS family of ∼5781 bp; this IS occurred twice at 2 different locations (with >99% sequence similarity, 100% coverage) in the chromosome of all our *E. faecalis* isolates (Table 2 and Suppl. Table 1). The novel transposable element harbored genes involved in its integration and excision, including genes for integrase, excisionase, replicase, a helix-turn-helix transcriptional regulator, methylated-DNA-protein-cysteine methyltransferase (MGMT), and trimethoprim-resistant dihydrofolate reductase (*dfrF*), which is involved in the resistance to diaminopyrimidine antibiotics (Figure 1D). The number of cargo genes and the length of this novel element suggest it is a unit transposon rather than an IS element. Unit transposons are larger than IS elements, as they may carry several passenger genes, usually ARGs, bound by inverted repeats (16, 48). Sequence similarity analysis of the entire region of 5781 bp against the NCBI database showed that this novel TE is 100% preserved and exists in multiple copies in the genomes of other *E. faecalis* strains, including 28157_4, 27725_1 and 27688_1. These *E. faecalis* genomes were sequenced using long and short reads and assembled following a hybrid *de novo* strategy (NCBI BioProject: PRJEB40976 from the University Medical Center of Utrecht). This transposable element could therefore be more widespread in *E. faecalis*, but its existence might have been concealed due to the lack of high-quality and complete assemblies for this species. According to CARD Prevalence 3.0.8 data, which was based on sequence data acquired from NCBI on August 13, 2020, the prevalence of the *dfrF* ARG in *E. faecalis* was low, being found in only 4.03% of all whole-genome sequences in this database (818 *E. faecalis* genomes). However, the prevalence of this gene in *E. faecium* species was 27.59% (1468 *E. faecium* genomes). Indeed *E. faecium* isolates from our study harbored several copies of *dfrF*. Searches of the novel transposable element family found in our *E. faecalis* isolates against the *E. faecium* isolates of our study resulted in *dfrF-dfrF* gene alignments with >99% sequence similarity and 100% coverage, but this was not the case for the other genes in the described transposable element from *E. faecalis* isolates. This novel element was the only IS found in the genetic neighborhood of ARGs in the chromosome of *E. faecalis* isolates.

We observed a different scenario when analyzing the genetic neighborhood of ARG in our *E. faecium* isolates, where 20 out of 21 ARG had at least one IS in their proximity (±10 annotated ORFs). We also observed that IS elements found in our *E. faecium* isolates were generally arranged as arrays. IS arrays consist of agglomerations of ISs in which several copies from the same IS family are proximal to each other, suggesting a high level of replicative transposition (49). Regarding their relationship with ARGs, we observed that the MFS transporter permease gene *efmA* was the only ARG in *E. faecium* that did not have IS elements in its proximity (Suppl. Table 1).

Differences in IS copy number between our 3 *E. faecium* isolates (Table 2) were directly related to reorganizations in modular plasmids caused by IS-mediated deletions in the form of composite transposons (Figure 1A, 1E). The conjugative plasmid pUAMSEF1a is a RepA_N megaplasmid (>260 kb) with a strikingly high number of IS sequences (n=59) (Table 2, Figure 1E) that was present only in the first *E. faecium* isolate we collected, UAMS_EF55. Despite the high number of IS families identified in this plasmid, only one ARG (aminoglycoside acetyltransferase, *aacA-aphD*) was found in pUAMSEF1a. However, there was a high number of efflux pumps, transporters, and phosphotransferase systems, along with several genes related to plasmid maintenance, replication, and transfer such as several toprim genes; DNA primases, topoisomerases, and resolvases; conjugal proteins; *tra* genes; and Type IV secretion system components. Although pUAMSEF1a was only identified in the UAMS_EF55 isolate, UAMS_EF57 and UAMS_EF58 harbored 2 smaller plasmids (pUAMSEF1b and pUAMSEF3), which resulted from the excision of pUAMSEF1a (Figure 1E, Suppl. Figure 2). The smaller plasmid product of pUAMSEF1a excision consisted of the segment bounded by 2 ISs in the same orientation, from the IS6 family in pUAMEF1a (specifically, 2 IS*1297* elements, first described in *Leuconostoc mesenteroides subsp. dextranicum* NZDRI2218). This circularized section constituted in its entirety the replicon pUAMSEF3 present in the 2 other *E. faecium* isolates (UAMS_EF57 and UAMS_EF58) that were collected just one and 2 days later, respectively, showing the high genetic activity of *E. faecium* isolates in this background. One of the 2 flanking IS6 elements was preserved in pUAMSEF1b (Suppl. Figure 3) after excision. However, both flanking IS6 elements were lost in the smaller plasmid pUAMSEF3 after the IS-mediated excision event. The plasmid pUAMSEF1b, found in UAMS_EF57 and UAMS_EF58 isolates, consisted of a RepA_N replicon of ∼205400 bp which is the largest circularized segment remaining after the IS-mediated excision of pUAMSEF3 (orange inner ring in Figure 1E). It is worth noting that another IS-mediated excision event was identified in the vancomycin resistance plasmid pUAMSEF4 in our *E. faecium* isolates; this is described in the section dedicated to the vancomycin plasmids.

### Daptomycin resistance mutations in *liaSR* genes are associated with a putative composite transposon in *Enterococcus faecium* isolates

Among the ISs inferred in the previous section, some ISs in the vicinity of ARGs in *E. faecium* were identified as potential composite transposons (note that experimental evidence for transposition of these IS-bounded structures has not been proven in this work). A noteworthy example of a putative composite transposon is the one found in the 3 *E. faecium* isolates formed by 2 IS*L3* elements identically oriented surrounding *liaSR* genes. LiaFSR is a three-component regulatory system involved in cell envelope homeostasis. As it has been reported in multiple studies (50, 51), amino acid substitutions in LiaR (W73C) and LiaS (T120A) are associated with a daptomycin (DAP)-resistant phenotype in *E. faecium*. Notably, all of our *E. faecium* isolates carried the LiaR^W73C^ and LiaS^T120A^ mutations. To test the hypothesis that there is an association between this IS*L3*-bounded structure and the presence of mutations associated with DAP resistance in *liaSR*, we extended our investigation to all complete, publicly accessible *E. faecium* genomes. Among 249 complete *E. faecium* genomes, we observed that 33 genomes carried both LiaR^W73C^ and LiaS^T120A^ mutations. Consistently, all 33 genomes also contained the same IS*L3*-bounded structure found in our *E. faecium* isolates (Figure 2A). In detail, this putative composite transposon is composed of 2 IS*L3* sequences identically oriented surrounding other 11 genes in almost all cases (Suppl. Figure 4). The genes flanked by the IS*L3* elements were identified as *tagD*, encoding a glycerol-3-phosphate cytidylyltransferase, which is central to the synthesis of teichoic acids in the cell wall of Gram-positive bacteria (52); an O-antigen ligase family protein gene involved in lipopolysaccharide biosynthesis (53); *wcaG*, encoding an epimerase; a serine hydrolase gene; *mltG*, encoding an inner membrane endolytic transglycosylase capable of cleaving at internal positions within a glycan polymer (54); *greA*, encoding a transcription elongation factor involved in bacterial invasion in other genera (55); a cell wall-active antibiotic-response protein gene; *liaS* and *liaR*; *trkA-C*, encoding a potassium-uptake protein; and *hesB*, encoding an iron–sulfur cluster biosynthesis protein.

**Figure 2.**
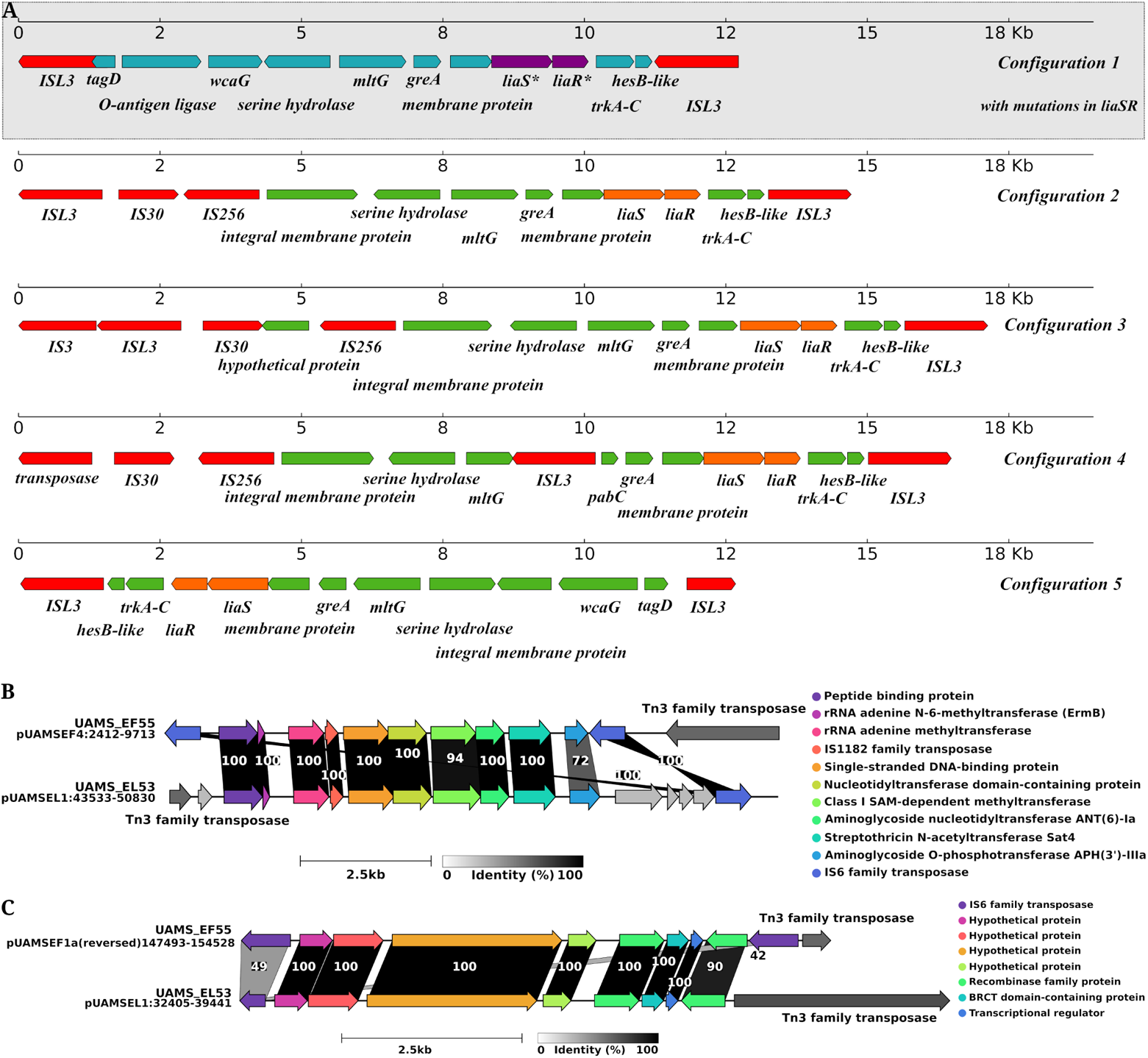
**A.** IS configurations surrounding *liaSR* genes in 252 complete *E. faecium* genomes. Configuration 1, highlighted in grey, was only found in *E. faecium* with mutations in *liaSR* genes (LiaS^T120A^ and LiaR^W73C^). **B.** Putative IS-mediated recombination event between *E. faecalis* and *E. faecium* strains with *emrB* genes and linkage genes *aad(6)-sat4-aph(3’)-IIIa.* **C.** Putative IS-mediated recombination event between pUAMSEF1a and pUAMSEL1 from *E. faecalis* and *E. faecium* isolates.

As observed previously (24, 56), the presence of LiaR^W73C^ and LiaS^T120A^ mutations does not always result in a DAP-resistant phenotype, but it has been related directly to poor bactericidal activity and further development of a DAP-resistant phenotype when DAP therapy was continued (24,57,58). To confirm that the arrangement of the putative composite transposition could be associated with the presence of LiaR^W73C^ and LiaS^T120A^ substitutions in *E. faecium* genomes, we surveyed a region of ∼15 000 bp that included *liaSR* genes in the other 217 complete *E. faecium* genomes that did not carry *liaSR* mutations. Among the strains with at least two ISs in the area surrounding *liaSR* genes, we identified 4 main IS-associated gene arrangements (Figure 2A). The most prevalent arrangement was “configuration 2,” which was present in 26 out of 249 surveyed genomes, followed by “configuration 3” found in 5 of the genomes. Most of the configurations contained variations in the copy number of some of the annotated ISs (mainly of IS*256* in “configuration 2”, which was duplicated in some strains). However, none of the configurations exhibited the “pseudo-compound transposon”–like structure found in association with the mutated *liaSR* genes in our isolates, with identically oriented IS elements from the IS*L3* family.

### *E. faecalis* and *E. faecium* isolates harbor 2 regions with IS-mediated recombination events

Sequence similarity analysis of all ISs found in our 6 isolates showed that the DNA sequence of some ISs (n=21) was highly conserved between the 6 isolates of this study, and those conserved ISs were found in several copies in different genomic regions (>99% sequence similarity, 100% coverage including IRs, transposase(s), and cargo gene(s) when applicable) (Suppl. Table 2). These highly conserved interspecific ISs (IS*3*, IS*6*, IS*256*, IS*982*, and IS*1182* families - elements IS*1485*, IS*1216*, IS*256*, IS*Efa13*, ISE*fm1*, and IS*1182*, respectively) belong to 5 families out of the 11 and 12 families found in *E. faecalis* and *E. faecium*, respectively. To investigate whether any of these highly conserved ISs had been transferred between our isolates of different species through IS-mediated recombination events, we extended the sequence similarity analysis to the regions flanking these highly preserved ISs by up to 4000 bp upstream and downstream from matched ISs between species. Our analysis identified 2 ∼7000 bp regions sharing >99.9% sequence similarity with 100% coverage between replicons from different species and bounded by IS elements (Figure 2B and 2C and Suppl. Table 2). Figure 2B depicts a putative IS-mediated recombination event between pUMASEL1 and pUAMSEF4 plasmids from *E. faecalis* and *E. faecium* isolates, respectively. Further investigation of this recombination event showed a high resemblance to a Tn5405-like element, mainly in *E. faecium* isolates. The Tn5405 element is a ∼12 kb composite transposon that was previously identified in methicillin-resistant staphylococci (59) and later described in *E. faecium* species (60). The gene clusters identified in our isolates contained the inducible by erythromycin *ermB* gene, 2 aminoglycoside ARG (*aad*(6) and *aph*(3’)-*III*a), and the streptothricin-resistance determinant gene *sat-4* (previously described linkage *aad(6)-sat4-aph(3’)-IIIa*) (60). The genes in the Tn5405 element are usually flanked by inverted repeats of 2 IS*1182* elements. In our isolates, only one IS*1182* was preserved inside the region, while the entire area was bounded by IS*6* family elements (specifically IS*1216*) on one side and by Tn3 family transposases (TnBth1 in case of *E. faecalis* and Tn4430 for *E. faecium* isolates) on the other side, which did not share sequence similarity (Figure 2B).

Immediately downstream of the region described above in pUAMSEL1 we found a second region of 7032 bp that was shared by all 6 isolates and was located in the pUAMSEF1a plasmid in *E. faecium* strains (Figure 2C). No ARGs were identified in this region, which harbored mainly a set of genes encoding hypothetical proteins along with a recombinase, a BRCT domain-containing protein (usually belonging to proteins involved in cell cycle checkpoint functions that respond to DNA damage), and a transcriptional regulator. These regions were bounded in both species by exactly the same elements as those found in the previously described event: 2 identically oriented IS*6* family elements (IS*1216*) and a Tn3 family transposase in one of the flanks in *E. faecium* isolates and one IS*6* family (IS*1216*) and a Tn3 family transposase in the case of *E. faecalis* isolates.

### Novel arrangement of IS elements in Tn1546-like transposon in *E. faecalis* isolates likely implicated in genotype–phenotype discrepancy

All of our *E. faecalis* isolates presented vancomycin MIC values of approximately 2 µg/ml, indicating they were vancomycin sensitive. However, all of these isolates also contained plasmids with complete *vanA* gene clusters known to confer vancomycin resistance, indicating a genotype–phenotype discrepancy (Figure 3). Between species, these *vanA*-containing plasmids differed in size, genetic content, and type of plasmid replication protein (*rep*) gene. However, the genetic sequences of all 7 genes in the *vanA* cluster (*vanR*, *vanS*, *vanH*, *vanA*, *vanX*, *vanY*, and *vanZ*) were highly conserved when aligned between isolates from both species (inner orange ring, Figure 3A and 3B).

**Figure 3.**
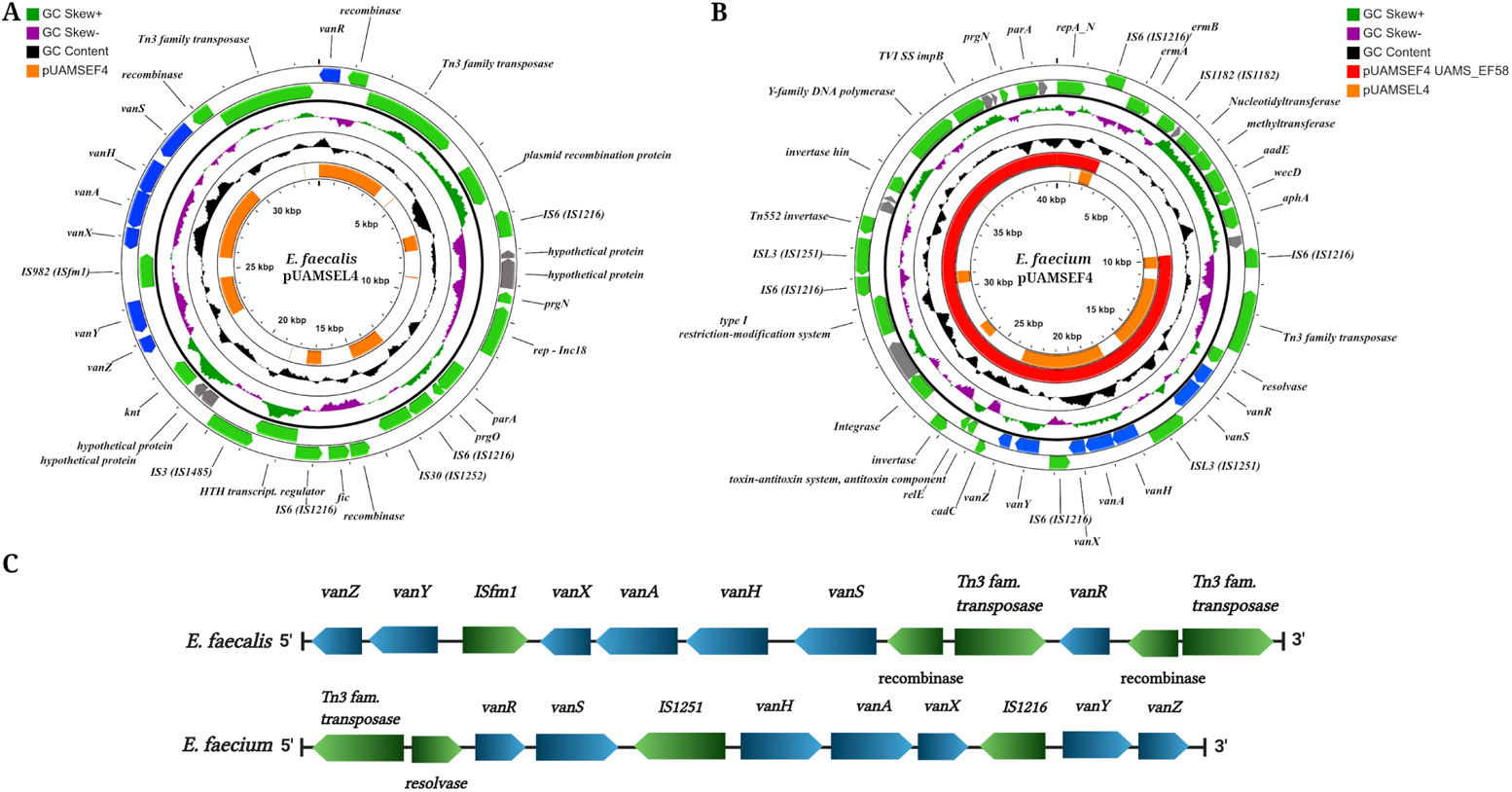
**A.** Full representation and functional annotation of the circularized vancomycin plasmid pUAMSEL4 from *E. faecalis*. Orange inner ring represents Blastn alignment against pUAMSEF4 from *E. faecium* UAMS_EF55. Width of the orange bar represents the percentage of sequence similarity (full width implies values closer to 100% sequence similarity). Genes annotated as hypothetical protein, are colored in grey. **B.** Full representation and functional annotation of the circularized vancomycin plasmid pUAMSEF4 from *E. faecium*. Orange inner ring represents Blastn alignment against pUAMSEL4 from *E. fecalis*. Red inner ring represents Blastn alignment against pUAMSEF4 from UAMS_EF58 isolate. **C.** Representation of *vanA* clusters and arrangements of ISs in *E. faecalis* and *E. faecium* isolates. Figures A and B were created using CGview server (version beta). Figure C was created using BioRender.

Plasmid pUAMSEL4 (carrying *vanA* gene cluster) from *E. faecalis* belongs to the incompatibility (Inc) 18-like family (61), which is a broad-host-range of conjugative plasmids that, under laboratory conditions, have been transferred successfully to a wide variety of microbes, such as *lactococci* (62), *streptococci* (63), and *staphylococci* (64, 65). On the other hand, the *E. faecium vanA* plasmid pUAMSEF4 was classified as belonging to the RepA_N family, which have a narrow host range (66). All *E. faecalis vanA* replicons were nearly identical in content and size, but we found a significant difference in the *vanA* plasmid from the last collected *E. faecium* isolate (UAMS_EF58) (Figure 3 and Suppl. Figure 1). This decrease in size in the UAMS_EF58 *vanA* plasmid was due to an IS-mediated deletion of a region located between the 2 identically oriented IS*6* elements (specifically, IS*1216)* located in the first quarter of pUAMSEF4 (red inner ring in Figure 3B, Suppl. Figure 5). Thus, pUAMSEF4 from UAMS_EF58 consisted of a replicon ∼7000 bp shorter than the *vanA* plasmids found in previously collected *E. faecium* isolates UAMS_EF55 and UAMS_EF57. In this excision event, both identically oriented IS*6* elements were retained in the *van* plasmid from UAMS_EF58, but none of the genes bounded by them were retained (Figure 3B, Suppl. Figure 5). The cluster of genes absent in pUAMSEF4 from UAMS_EF58 contained 7 ARGs that code for the resistance to aminoglycosides (*aad*(*6*)*, aph(3’)-IIIa*), nucleosides (*sat-4*) and macrolides, lincosamides and streptogramins (*ermAB*) (Figure 3B).

Although the sequence of genes in the *vanA* cluster was highly preserved between our 6 isolates, *vanA* gene clusters from both species presented different organizations of the Tn1546-like element. The Tn1546-like transposon in pUAMSEF4 from *E. faecium* isolates (Figure 3B) showed a genetic organization that is highly preserved between hospital-acquired vancomycin-resistant *E. faecium* isolates (67). However, we observed an unusual genetic organization of the Tn1546-like transposon in our *E. faecalis* isolates; this peculiar organization of ISs in the *vanA* cluster was not found in any other strain when the entire region was queried in the NCBI public database (Suppl. Figure 6). It has been previously reported that the *vanA* cluster (*vanRSHAXYZ*) is prone to IS-mediated alterations that can affect the resistant phenotype of a strain, even in the case of a *vanA*-positive genotype (68, 69). We speculate that this novel arrangement of IS elements in the Tn1546-like transposon in *E. faecalis* isolates with a *vanA*-positive genotype from our study might be the cause of the “silenced” phenotype observed in the experimental analysis performed when collecting these isolates.

## Discussion

Enterococci are widely distributed across many different niches, but their primary habitat is the gastrointestinal tract (70). Among enterococci, *E. faecalis* is the most common cause of opportunistic infection, but *E. faecium* is intrinsically more resistant to antibiotics. More than half of nosocomial *E. faecium* isolates in the US have expressed resistance to ampicillin and vancomycin and high-level resistance to aminoglycosides (71). While the relationship between ISs and ARGs has been established (16,18,48), there is lack of information on the association between ISs and ARGs and other aspects regarding the nature, maintenance, and mechanisms of transmission of ISs in bacterial genomes.

Although mutations produced by ISs are generally deleterious (46, 72) and bacterial genomes possess mechanisms that suppress transposition (72–74), a high level of transposition activity has been understood as the mechanism of adaptation to stressful and dynamic environmental conditions that can generate sufficient genetic variation to increase fitness (75). In this work, we present 2 bacterial species sharing the same environment (patient’s bloodstream infection) with a very similar content of IS families but drastically different copy numbers and arrangement of ISs throughout their replicons. *E. faecium* genomes harbored a strikingly high number of ISs and showed a higher level of IS-mediated activity, even in a short period of time (a total of 4 days between first and last sample collected. Considering the first isolate collected for each species as the reference (UAMS_EL53 for *E. faecalis* and UAMS_EF55 *E. faecium*), the only significant genomic differences found between the isolates of the study were the IS-mediated events, highlighting the significant role of ISs and other transposable elements as mechanisms that rapidly generate adaptation in bacterial genomes (18). The high number of ISs found in our isolates compared to previous reports (76) could be due partially to our use of high-quality, complete genomes, considering draft assemblies are typically highly fragmented and their contigs are usually broken at IS locations (77) due to the repetitive nature of IS elements. In agreement with previous reports, our 6 isolates showed a strong correlation between IS elements and ARGs in the sense that the majority of ARGs had at least one IS element in their surrounding regions, suggesting that these IS elements might have a role in the transfer of these ARGs and other virulence factors and determinants of antibiotic resistance (16, 48).

The presence of a high number of IS families with highly conserved DNA sequences between *E. faecalis* and *E. faecium* strains suggests that these elements could have been spread through horizontal gene transfer. Most importantly, IS propagation generates multiple homologous sequences scattered throughout the bacterial genomes, which are substrates for homologous recombination (49), paving the way for a successful transmission of the antibiotic-resistance determinants and virulence factors that frequently accompany these elements in *E. faecalis* and *E. faecium* genomes.

It has been documented that elements from the IS*6* family with flanking ISs in direct repeat are able to form transposons that resemble compound transposons through a mechanism that involves the formation of cointegrates, also known as pseudo-compound transposons (78,79,80), and also via circular translocatable units (80, 81). Potential mechanisms of circular molecular transposition (translocatable units) mediated by elements from IS*6* and IS*26* families have been described (80, 82). Transposon-derived circular forms have been documented in Gram-negative bacteria such as *Acinetobacter baumannii* (83) and *Escherichia coli* (84). In this work, we provide evidence of the formation of a complete conjugative plasmid (pUAMSEF3) with all of the necessary machinery and functional elements for its own replication and transference (Figure 1F); this plasmid was the product of IS-mediated excision from a larger plasmid (pUAMSEF1a) (Figure 1E), supporting the key role of translocatable units in disseminating genetic material as autonomous entities.

We also observed multiple putative pseudo-compound transposons when analyzing the genomes of our 3 *E. faecium* isolates. However, this type of structure had never been described before, to the best of our knowledge, for members of the IS*L3* family. The novel structure is composed of 2 directly oriented IS*L3* elements surrounding *liaSR* with daptomycin-resistant associated mutations and surrounding genes. The extended analysis to all complete, publicly available *E. faecium* genomes confirmed that the novel structure is specific to *E. faecium* genomes with daptomycin-resistance mutations in *liaSR* genes. Our findings suggest a new paradigm for the investigation of daptomycin-resistance mechanisms and their dissemination in *faecium* species, in the sense that the development of a daptomycin-resistant phenotype might not be produced by sequential co-mutations in *liaSR* genes but through IS-mediated transmission of the entire set of genes already carrying these mutations. However, we found no duplication of this cluster in our *E. faecium* isolates or in the other 33 complete *E. faecium* genomes harboring *liaSR* mutations. In the same vein, the upstream and downstream genetic neighborhood of the IS*L3* bounded structure, was shared between all complete isolates queried, therefore homologous recombination might be a more plausible mechanism of propagation for these mutations in *liaSR* genes than ISs mediated transposition. Homologous recombination mediated by ISs between related strains has been previously documented in other works (85–87). Further detailed examination of the ISs duplications at target sites and experimental assays should be carried out to determine if this ISL3 bounded structure, observed in all *E. faecium* genomes found carrying amino acid substitutions in LiaR (W73C) and LiaS (T120A), is crucial in their dissemination. Our *E. faecalis* isolates harbored a complete *vanA* cluster but presented MIC values ≤ 2 for vancomycin, indicating genotypic–phenotypic discrepancy. Among the most widely documented effects of IS transposition is gene inactivation (18, 88), and the novel arrangement of ISs found in the *vanA* cluster of our 3 *E. faecalis* isolates might explain the observed phenotypic discrepancy. Similar events have been documented in other species (68, 69). A particularly interesting case was described by Siversten and colleagues (69) in which initially vancomycin-susceptible *E. faecium* isolates were able to survive for several days during subclinical breakpoint exposure to glycopeptides until they developed the resistance phenotype after the excision of one IS element present in the *vanA* gene cluster.

In summary, while our computational results are all biological observations that need to be explored experimentally, our results substantiate that IS-mediated events differ between 2 related species such as *E. faecium* and *E. faecalis* when sharing the same environment and under complex selective forces occasioned by intensive antibiotic treatment. In addition, some of these IS-mediated reorganizations are noticeable even between isolates collected from the same subject in a short period of time. Our results highlight the major role of ISs and other transposable elements in the rapid adaptation and response of enterococci to clinically relevant stresses.

## Data availability

Raw sequencing data, genome assemblies, and functional annotations for the isolates described in this study are available under the BioProject accession number PRJNA735268 in the NCBI database.

## Author contributions

SJ and AK conceived of the study. SJ and ZU designed the study. AK reviewed clinical metadata. ZU performed the bioinformatics analysis. ZU drafted the manuscript. ZU and SJ edited the manuscript. KA assisted with insertion sequence distribution analysis and Figure 1 creation. SJ and ZU were responsible for data interpretation. All authors have read and approved the final manuscript. Editorial support was provided by the Science Communication Group at the University of Arkansas for Medical Sciences.

## Ethics Declarations

### Ethics approval and consent to participate

This study was approved by the Institutional Review Board of the University of Arkansas for Medical Sciences (IRB No. 228137).

## Supplementary Files

**Suppl. Table 1.** Annotation tables of the 6 isolates. The functional annotation was performed using Prokka (35), Antimicrobial resistance (AMR) determinants by RGI from the CARD (36), and IS families annotation by ISEScan (37) were included and sorted by replicon coordinates.

**Suppl. Table 2.** Blastn output of ISs alignments filtered by sequence similarity and sequence coverage.

**Suppl. Figure 1.** Genomic tree of Mash distances using the DNA sequence of all plasmids in this study. Mash distances between the complete genomes utilized in this study are represented in the table showed beneath the tree.

**Suppl. Figure 2.** Overview of the alignment of the Oxford Nanopore reads from UAMS_EF58 against pUAMSEF1a plasmid from UAMS_EF55. Figure points out the area of the IS-mediated excision events that generated pUAMSEF3 in UAMSEF_57 and UAMS_EF58. In the central region of the figure, the discontinuity produced by the reads from pUAMSEF3 inserted in pUAMSEF1a can be observed (black arrows). Coverage of Oxford Nanopore reads in this area was >400x.

**Suppl. Figure 3.** Circular representation of pUAMSEF1b plasmid from UAMSEF_57 and UAMSEF_58 isolates.

**Suppl. Figure 4.** Representation of clusters of genes from 33 complete *E. faecium* genomes that harbor genes with mutations in *liaSR* genes (LiaS^T120A^, LiaR^W73C^) and surrounded by IS*L3* elements. Numbers below coding genes represent sequence similarity in percentage between the orthologs in the cluster. The similarity is not shown if two orthologs have sequence similarity < 0.5 (50%). Genes sharing same color have same functional annotation and have high level of sequence similarity. Genes without orthologs in this figure, were represented in grey color. **Suppl. Figure 5.** Overview of the alignment of the Oxford Nanopore reads from UAMS_EF58 against pUAMSEF4 plasmid from UAMS_EF55. Figure points out the area of the IS-mediated excision event that is deleted in pUAMSEF4 plasmid from UAMSEF_58 isolate. Note that the lowest coverage in that area was >600x.

**Suppl. Figure 6.** Overview of the alignment of the *vanA* cluster from *E. faecalis* isolates against the Blastn database. The *vanA* cluster from our isolates, did not completely align with any of the genomes from the nr/nt nucleotide collection from NCBI. Higher similarities were found against the *E. saigonensis vanA* cluster.

