## Supplementary figures and images for "Insertion sequences associated with antibiotic resistance genes in *Enterococcus* isolates from an inpatient with prolonged bacteremia"

### Supplementary Figure 1

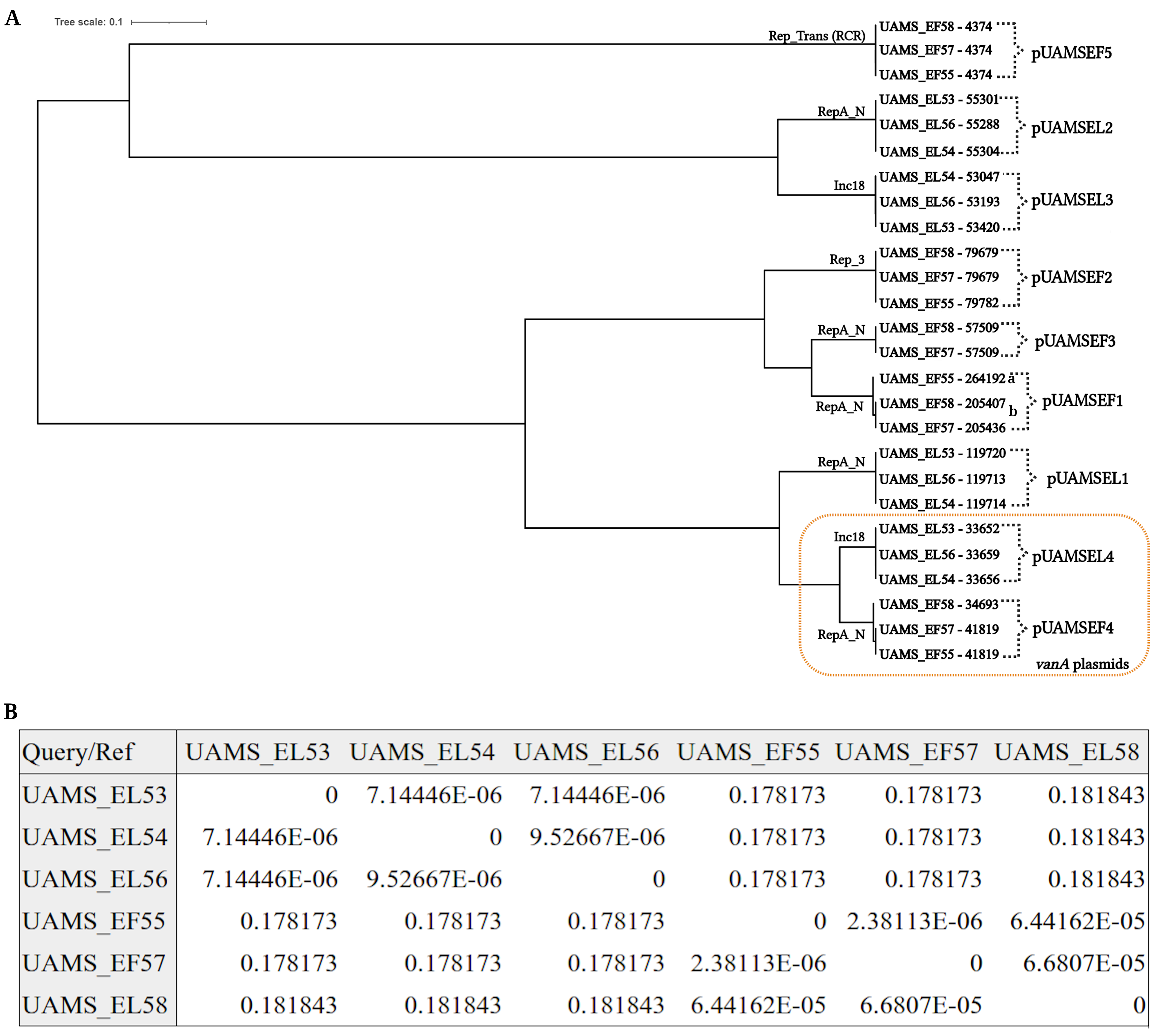

### Supplementary Figure 2

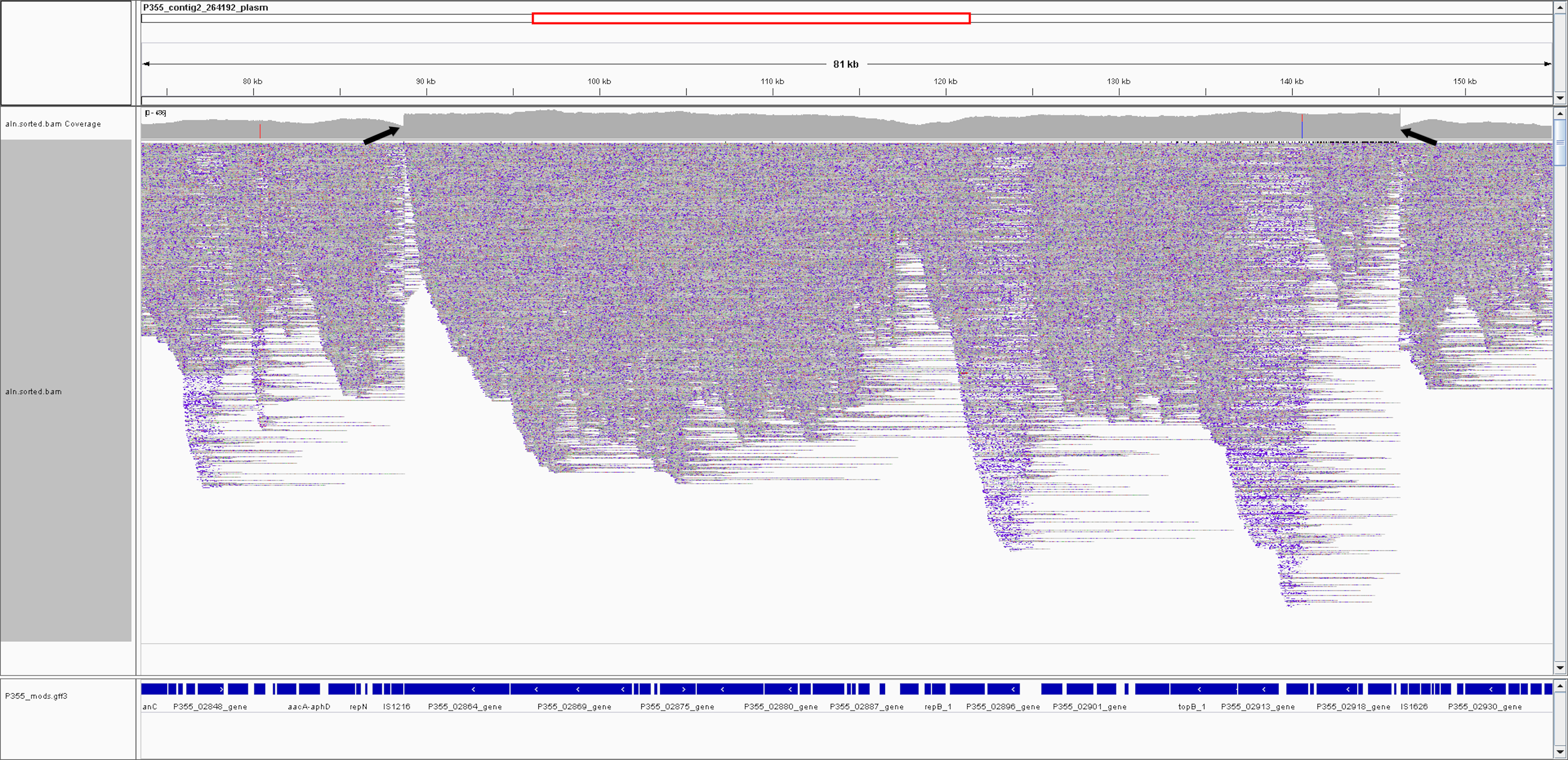

### Supplementary Figure 3

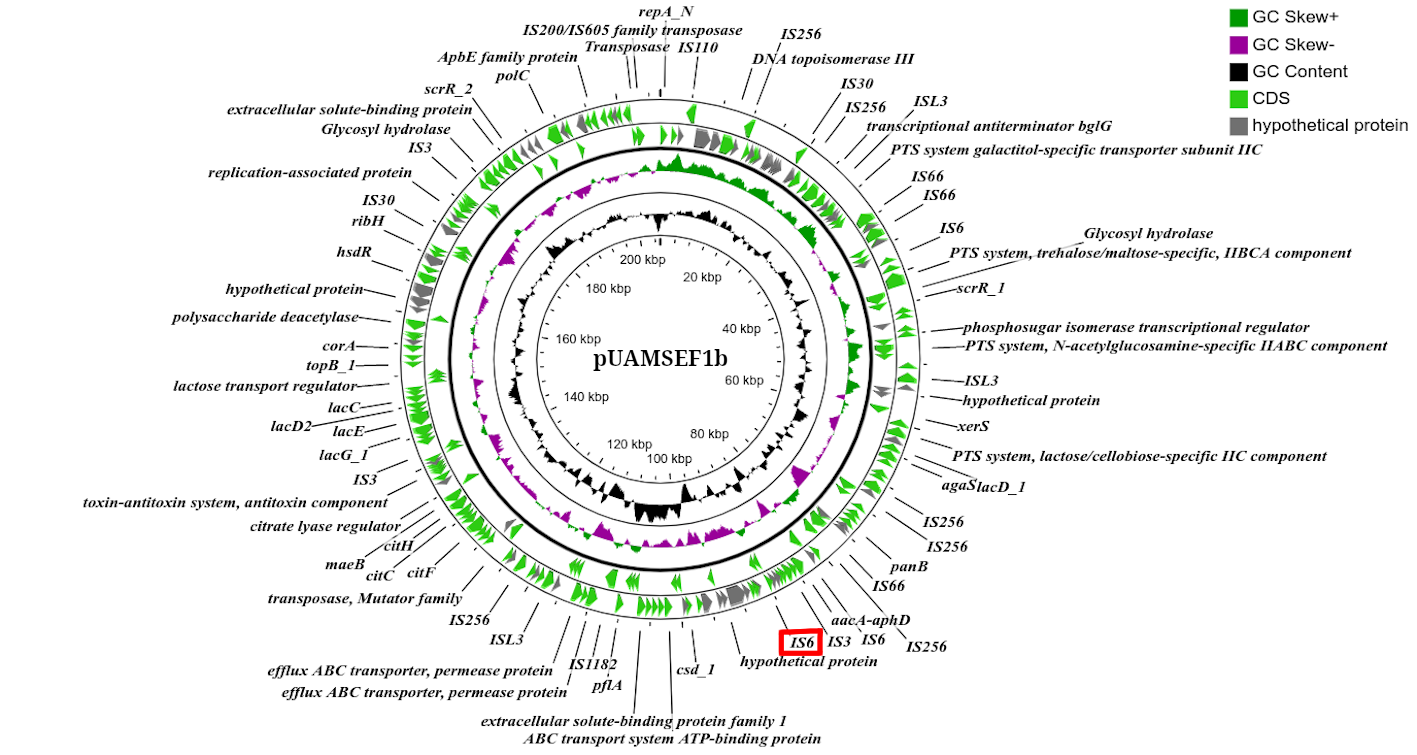

### Supplementary Figure 4

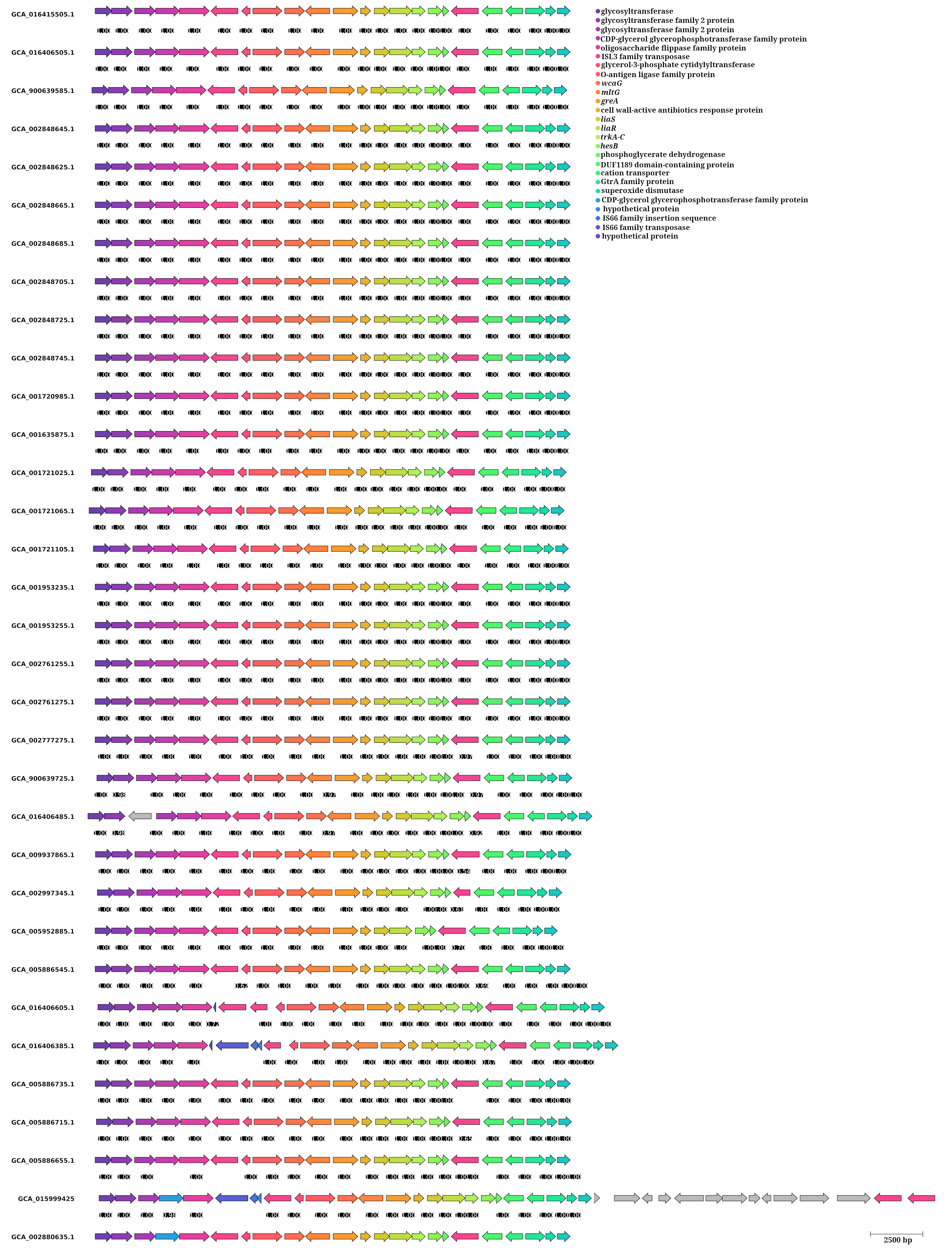

### Supplementary Figure 5

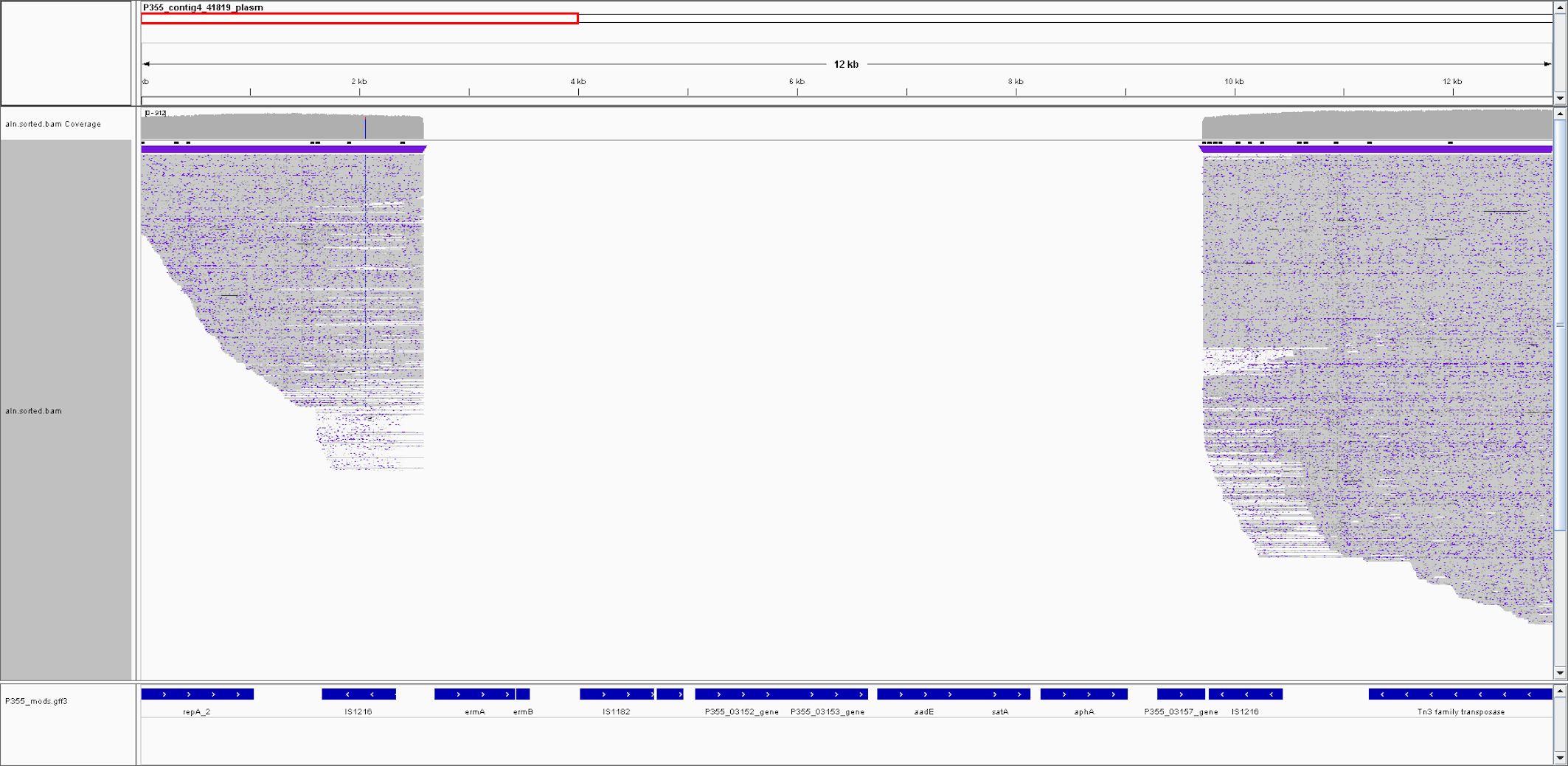

### Supplementary Figure 6

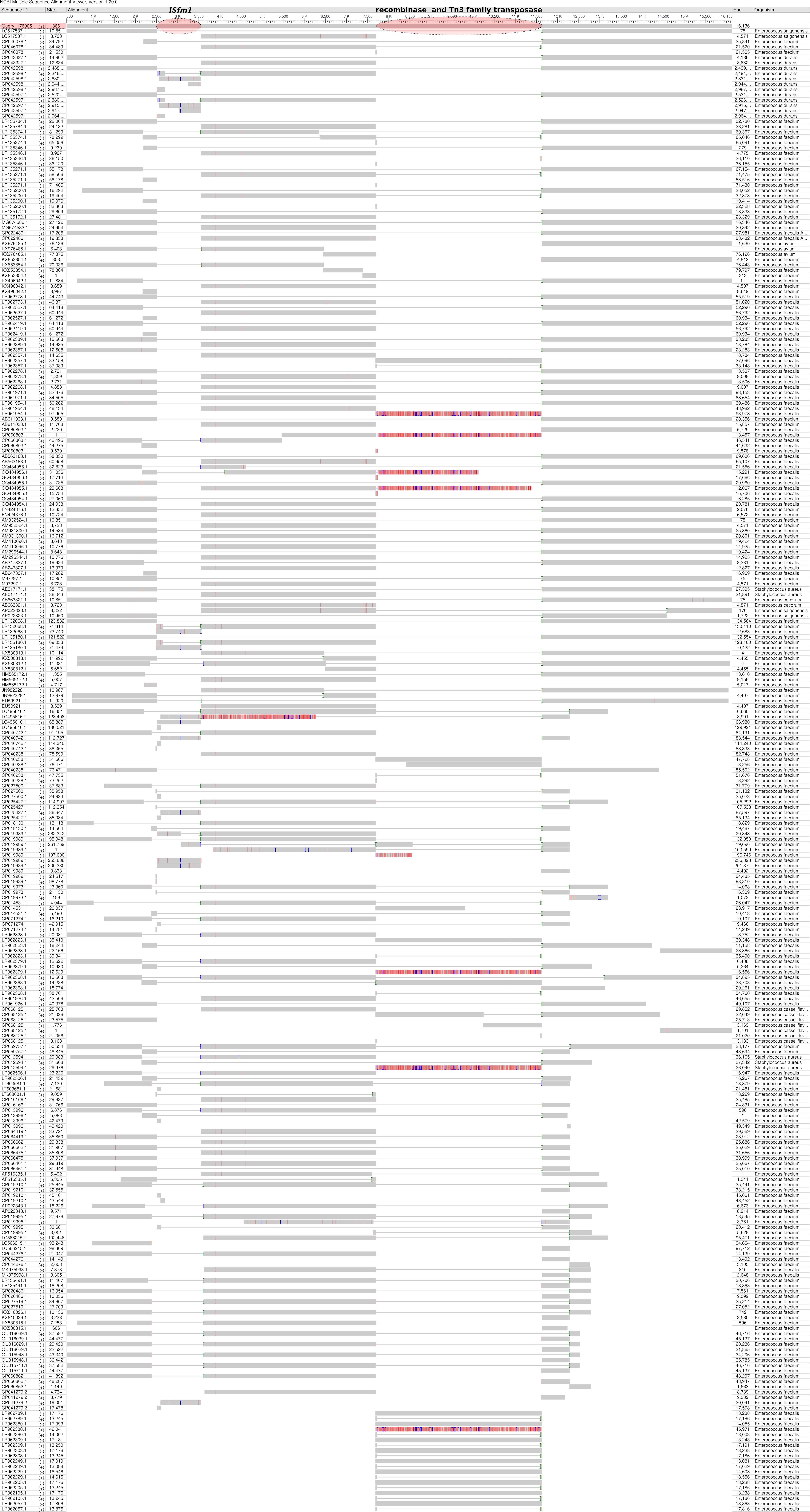
